# Arginine methylation of BAF155 regulates interactions with RNA processing machinery

**DOI:** 10.64898/2026.05.18.726059

**Authors:** Mallory Sokolowski, Deena Scoville, Peyton C. Kuhlers, Jesse R. Raab

**Author notes:** **Correspondence**: Jesse. R. Raab, Department of Genetics, Lineberger Comprehensive Cancer, Center University of North Carolina at Chapel Hill.

## Abstract

Post-translational modifications (PTMs) of chromatin remodelers are abundant but functionally understudied. Here we investigate the role of asymmetric dimethylation of arginine 1064 (BAF155me2a) on the SWI/SNF core subunit BAF155, a mark deposited by CARM1/PRMT4 that has been linked to tumor progression but whose molecular function remains unclear. Using immunoprecipitation–mass spectrometry with a dimethyl-specific antibody, we found that R1064me2 selectively enhances BAF155 interactions with RNA processing factors, including the anti-termination protein SCAF4, splicing factors, and the transcription factor RFX5. CUT&RUN profiling showed that BAF155me2a, SCAF4, and RFX5 co-occupy promoter regions, and reciprocal immunoprecipitations confirmed that the SCAF4–BAF155 interaction depends on R1064 methylation. To test the functional consequences of this modification, we generated cells expressing either wild-type BAF155 or a methylation-deficient BAF155-R1064K mutant. Loss of methylation did not alter chromatin accessibility, BAF155 genomic occupancy, or SCAF4 recruitment. However, nascent transcription measured by TT-seq revealed a coordinated reduction in 5′ sense transcripts and upstream antisense transcripts (PROMPTs) at BAF155-bound promoters, with a quantitatively larger decrease in PROMPTs at SCAF4 co-bound sites. The effect was restricted to the promoter-proximal region and resolved toward the gene end, consistent with a defect in productive elongation downstream of RNA polymerase II recruitment. These data support a model in which BAF155 dimethylation provides a co-transcriptional interface coupling SWI/SNF to RNA processing machinery, and identify regulation of nascent transcription as a non-canonical function of SWI/SNF PTMs.

## Introduction

Chromatin regulation and proper control of gene expression is crucial to organism survival and function. This system hinges upon key protein complexes that can recognize and bind chromatin and facilitate changes to ensure the correct timing of transcription and response to environmental cues [1,2]. The Switch/Sucrose Non-Fermentable (SWI/SNF) chromatin remodelers are a family of highly conserved protein complexes with an established role in regulating chromatin and coordinating gene expression [3,4]. SWI/SNF complexes have an ATPase module that allows for the use of ATP to efficiently remodel chromatin through the sliding and eviction of nucleosomes [5,6]. There are three main forms of the SWI/SNF complexes: canonical BAF (cBAF), polybromo BAF (PBAF), and non-canonical BAF/GLTSCR1/like-containing BAF complex (ncBAF/GBAF) [7–10]. Each subcomplex has complex specific subunits as well as core subunits that are shared [11]. Mutations in SWI/SNF complexes are found in ∼20% of all cancers [12] and loss of specific subunits is embryonic lethal in mice [13–15].

SWI/SNF complexes are able to modify different regions of chromatin through the recognition of post-translational modifications (PTMs) on histone tails [16,17]. This interaction leads to the recruitment of other cofactors to free or restrict the corresponding DNA wrapped around the surrounding histones. While recognizing these modifications, SWI/SNF complexes themselves can be post-translationally modified. A proteome-wide study of arginine monomethylation identified multiple methylation sites on several SWI/SNF subunits, among other chromatin remodeling complexes [18]. The ATPase subunit, BRG1/SMARCA4, is phosphorylated by casein kinase 2 (CK2) in primary murine myoblasts during mitosis to promote proliferation [19]. SWI/SNF complex integrity is regulated by LSD1 mediated demethylation of monomethylated lysine residues on BAF155/SMARCC1 and BAF170/SMARCC2 during development [20]. These observations suggest that PTMs on the SWI/SNF complex are important and may guide complex function.

Alongside monomethylation, asymmetric dimethylation of an arginine residue on the SWI/SNF subunit, BAF155, was observed in triple-negative breast cancer [21], which enhanced tumor growth and aggression. Modified BAF155 was required for tumor growth and invasion in this model through the recruitment of co-activator, BRD4, to promote oncogene expression via super-enhancers and repress immune responses [22], highlighting that PTMs on non-histone substrates have disease relevance. CARM1/PRMT4 [21] deposits the asymmetric dimethylation mark on arginine 1064 (R1064) in a disordered region near the C-terminus of BAF155 (Figure 1A). However, it remains unclear how BAF155 methylation impacts SWI/SNF complex formation and if this affects protein interactions.

**Figure 1.**
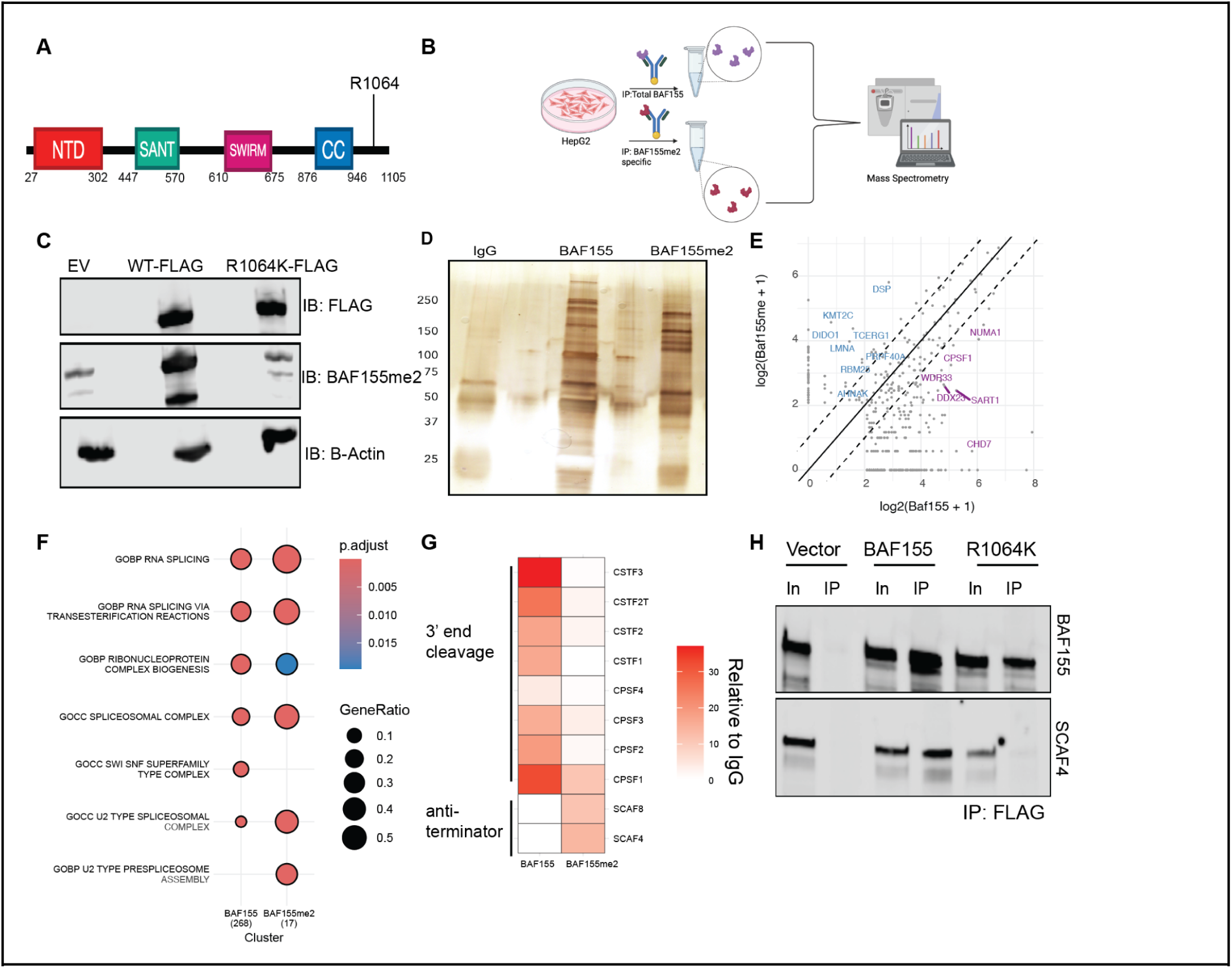
Asymmetric dimethylation on arginine 1064 of BAF155 influences protein-protein interactions. **A.** Schematic of BAF155 protein domains and location of dimethyl mark (NTD = N-terminal domain, SANT = Swi3, Ada2, N-Cor, and TFIIIB domain, SWIRM = Swi3p, Rsc8p and Moira domain, CC = coiled-coil domain). **B.** Schematic of immunoprecipitation-mass spectrometry (IP-MS) approach in HepG2 cells. **C.** HEK293T cells were transfected with the indicated epitope-tagged constructs. BAF155, BAF155me2a, and FLAG were visualized by western blot. **D.** Silver stain of resulting lysates of BAF155, BAF155me2a, and IgG (control) immunoprecipitations that were used in downstream mass-spectrometry. **E.** Log2 fold change (Log2FC) of proteins detected in immunoprecipitation-mass-spectrometry in the BAF155 lysate plotted against the Log2FC in the BAF155me2a lysate (p-value < 0.05). Proteins with a Log2FC of BAF155 relative to BAF155me2a between −1 to 1 are colored black, Log2FC > 1 is colored magenta, Log2FC < −1 is colored light blue. **F.** Dot plot of enriched Gene Ontology Biological Processes based on proteins found in either the BAF155 or BAF155me2a specific lysate from our IP-MS data. **G.** Heatmap of spectral counts for proteins involved in 3’ end mRNA processing in the BAF155 and BAF155me2a IP-MS. **H.** Immunoprecipitations for FLAG in HEK293Ts with the indicated overexpression constructs. Western blot analysis for SCAF4 and total BAF155 was performed in the input and resulting lysate for each cell line.

The abundance of PTMs on chromatin remodelers and the well-established relationship between the SWI/SNF complexes and cancer underline the importance of understanding the function of these modified complexes. The role of SWI/SNF complex PTMs is understudied but can provide functional insights into how these modifications drive phenotypic outcomes. We sought to understand the function of asymmetric dimethylation on BAF155 and the consequences of loss of this mark.

Here, we found that asymmetric dimethylation on BAF155 guides SWI/SNF protein-protein interactions and is enriched in RNA processing factors. We saw that dimethylated BAF155 targets promoters co-bound by RNA processing factors and the transcription factor, RFX5, to regulate gene expression. We observed that loss of R1064me2 results in a perturbed transcriptional profile at the 5’ and upstream antisense region of genes that is independent of chromatin accessibility and BAF155 targeting.

## Results

### Arginine dimethylation alters SWI/SNF complex interactions with RNA processing machinery

To investigate how BAF155 methylation influences SWI/SNF complex assembly and protein interactions, we performed immunoprecipitation-mass spectrometry (IP-MS) using either a total BAF155 antibody or an antibody that recognizes the asymmetric dimethyl specific form of BAF155 (BAF155me2a) in HepG2, a hepatoblastoma cell line (Figure 1B). To validate antibody specificity we transfected HEK293T cells with an epitope tagged WT or BAF155-R1064K mutant that cannot be methylated (Figure 1C). The R1064K point mutant prevents CARM1/PRMT4 from methylating BAF155 while maintaining a positive charge on BAF155. We performed these experiments under moderate stringency wash conditions (300mM NaCl) to recover proteins that may interact with the SWI/SNF complex but were not SWI/SNF complex members. A silver stain of the two resulting immunoprecipitations in HepG2 cells showed a marked difference in interacting proteins (Figure 1D). We then subjected these immunoprecipitations to mass spectrometry to identify novel interactions specific to asymmetrically dimethylated BAF155. Comparison of ncBAF/GBAF, PBAF, and cBAF subcomplex members as well as shared subunits in the total BAF155 antibody pull down to the dimethyl specific antibody revealed no difference or preference in SWI/SNF complex formation (Supplemental Fig. 1). The lack of change or complex preference suggests that the methylation status of BAF155 does not impact SWI/SNF complex assembly.

We broadened our analysis to all other interacting proteins and compared the enrichment in BAF155 total IPs relative to BAF155me2a IPs (Figure 1E, Supplemental Table 1). Performing Gene Set Enrichment Analysis (GSEA) on the proteins significantly enriched in either IP revealed a strong enrichment of RNA processing factors in both the total BAF155 IP and the dimethyl-specific IP (Figure 1F). Notably, we observed some highly specific interactions that were more strongly enriched in either total BAF155 IPs or dimethyl specific IPs. For example, we observed RNA binding proteins, anti-termination factors, such as SCAF4 and SCAF8, and splicing factors in the BAF155me2a-specific immunoprecipitation (Figure 1E, G). Conversely, polyadenylation and 3’ end cleavage machinery was enriched in the total BAF155 antibody immunoprecipitation along with other known BAF155 interactors, including CHD7 [23] (Figure 1E, G).

SCAF4 has been shown to be a key regulator of 3’ end site choice, but its association with chromatin remodeling factors has not been observed [24]. To validate that the interaction between BAF155 and SCAF4 is mediated by the methylation status of BAF155, we performed an IP for FLAG and immunoblotted for BAF155 and SCAF4 in our established HEK293T overexpression cell lines. We observed a distinct loss of SCAF4 interaction when BAF155 can no longer be methylated compared to the WT construct (Figure 1H). This loss suggests that the SCAF4-BAF155 interaction is dependent on the methylation status of R1064 and that methylation status of BAF155 is important in guiding its protein interactions with RNA processing machinery.

### BAF155 and SCAF4 are co-bound at promoter regions

We identified a previously unknown interaction between the anti-termination factor, SCAF4, and BAF155 which is mediated by the methylation status of R1064 on BAF155. To facilitate study of the function of this modification, we generated a liver cancer cell line model (HLF) by using CRISPR/Cas9 to knock out endogenous BAF155 (BAF155-KO) and stably re-expressed an epitope tagged (FLAG) BAF155-WT or BAF155-R1064K (Figure 2A). SCAF4 has previously been shown to interact with RNA, but chromatin binding has not been tested [24]. To further define the interaction of BAF155me2a and SCAF4 and broadly profile BAF155 binding when R1064me2 is disrupted, we performed CUT&RUN [25] in our cell line model. This approach allowed us to map the location of BAF155-FLAG, BAF155me2a, SCAF4 and the histone mark H3K27ac. In both BAF155-FLAG and BAF155-R1064K-FLAG, we observed widespread overlap of BAF155me2a and BAF155-FLAG CUT&RUN signals (Supplemental Figure 2A-2C). While prior ChIP-seq studies observed differences in BAF155me2a and total BAF localization, the higher specificity of CUT&RUN may explain the difference. While peak overlaps suggest some BAF155me2a-only or FLAG-only regions, this is likely due to peak calling thresholds as signal intensity is similar at these sites in both WT and BAF155-R1064K (Supplemental 2A-C) [21,22].

**Figure 2.**
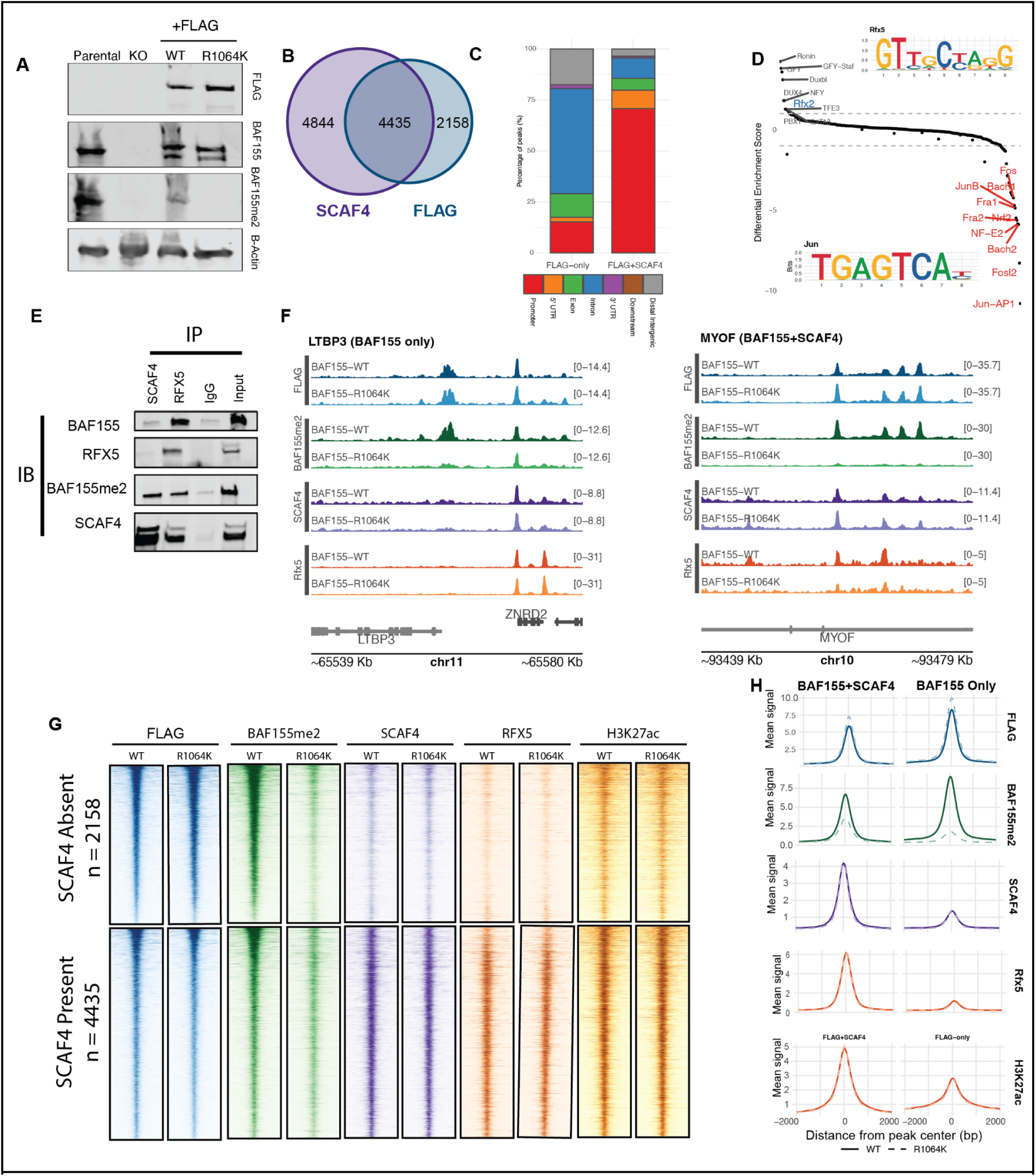
SCAF4-BAF155 interaction is driven by R1064 methylation and cooperatively bind promoters. **A.** Western blot verification for HLF cell lines that were engineered to have a BAF155 KO background and CRISPR knock-in for a WT version of BAF155 or a R1064K point mutant. The knock-in cell lines are also epitope tagged. **B.** Overlap of peaks for BAF155-FLAG and SCAF4 CUT&RUN in engineered HLF cell lines. (p-value < 2.2e-16, hypergeometric test) **C.** Proportion of BAF155-FLAG alone, SCAF4 alone, and overlapping peaks at enhancer and promoter regions. **D.** Differential score of HOMER motif analysis of peaks bound by BAF155-FLAG alone compared to motifs enriched at BAF155-FLAG and SCAF4 peak overlaps. RFX5 consensus binding motif identified through HOMER (upper right corner). **E.** Reciprocal IPs for SCAF4 and RFX5 and corresponding western blot for BAF155, BAF155me2a, SCAF4, and RFX5 in HepG2 cells. **F.** Genome browser examples of BAF155-FLAG CUT&RUN peaks bound independently and F. co-bound with SCAF4 and RFX5. **G.** Heatmaps of CUT&RUN signal for FLAG, BAF155me2a, SCAF4, RFX5, H3K4me3, and H3K27ac at regions bound by BAF155 alone (n = 2158) or co-bound by both BAF155 and SCAF4 (n = 4435). Signal is centered around the transcriptional start site (TSS). **H.** Metaplots of quantification in G.

We next compared SCAF4 and BAF155me2a occupancy across the genome. SCAF4 interacts with RNA, but direct genomic binding had not been previously shown [24]. CUT&RUN analysis revealed that ∼48% of SCAF4 bound regions overlap BAF155-FLAG binding sites (Figure 2B). BAF155+SCAF4 occupied regions were heavily biased towards promoters, while BAF155 lacking SCAF4 was more associated with intronic and distal intergenic regions (Figure 2C). We used BAF155-FLAG as a proxy for BAF155me2a occupancy given their considerable overlap. We separated genomic regions into those bound by both BAF155-FLAG and SCAF4 and those bound only by BAF155-FLAG and performed HOMER motif analysis. This revealed an enrichment for members of the RFX family as among the most preferentially found in BAF155+SCAF4 cobound regions (Figure 2D, Supplemental Figure 2D). Notably, RFX5 was enriched in our dimethyl specific IP-MS data, validating this association.

To further validate the interactions between BAF155, BAF155me2a, SCAF4, and RFX5, we performed multiple reciprocal IPs in HepG2 cells and immunoblotted for each protein respectively. We found that RFX5 could immunoprecipitate BAF155 and SCAF4 (Figure 2E). Moreover, BAF155me2a and SCAF4 showed a stronger interaction than total BAF155 when enriching for SCAF4 (Figure 2E). This is consistent with our IP-MS data and CUT&RUN analysis. RFX5 and SCAF4 displayed a distinct interaction with each other across IP conditions, supporting our previous observations of co-localization at promoter regions (Figure 2E).

To verify if RFX5 binds to the same regions of the genome as SCAF4 and BAF155me2a, we performed CUT&RUN for RFX5 in BAF155-WT and BAF155-R1064K cell lines. We observed co-localization of RFX5, SCAF4, and BAF155me2a at promoter regions (Figure 2F-H). Notably, we did not detect a global decrease in SCAF4 or RFX5 when BAF155 could not be methylated (Figure 2G,H). RFX5 was specifically co-localized to regions also containing SCAF4/BAF155, consistent with our IP data (Figure 2E, 2G-H). Our findings suggest that methylation of R1064 on BAF155 is associated with a complex of SCAF4 and RFX5 at promoters.

### BAF155me2a is dispensable for maintaining the chromatin landscape

To determine if loss of BAF155me2a would impact chromatin accessibility, we performed ATAC-seq and subsequent differential accessibility analysis in our established HLF cell lines. ATAC-seq revealed no statistically significant changes in chromatin accessibility when BAF155 cannot be methylated (Figure 3A). The majority of open chromatin sites (∼125,000 peaks) did not experience a change in accessibility when BAF155me2a or total BAF155 was lost and differential accessibility analysis did not identify any statistically significant differences (Figure 3A, B). The moderate differences in peak numbers and overlap are likely due to peak thresholding differences where ‘Shared’ peaks had much higher open chromatin signals (Supplemental Figure 3A, B). We compared open chromatin specifically at regions bound by SCAF4 and BAF155-FLAG compared to those bound only by BAF155. Similar to CUT&RUN, open chromatin at BAF155+SCAF4 bound regions were more associated with promoters, while BAF155 regions lacking SCAF4 were more enriched for distal intergenic and intronic regions (Figure 3C). Similar to the overall levels of ATAC signal changes, there were no differences in ATAC-signal between WT and R1064K mutants at BAF155 only regions or BAF155+SCAF regions (Figure 3D).

**Figure 3.**
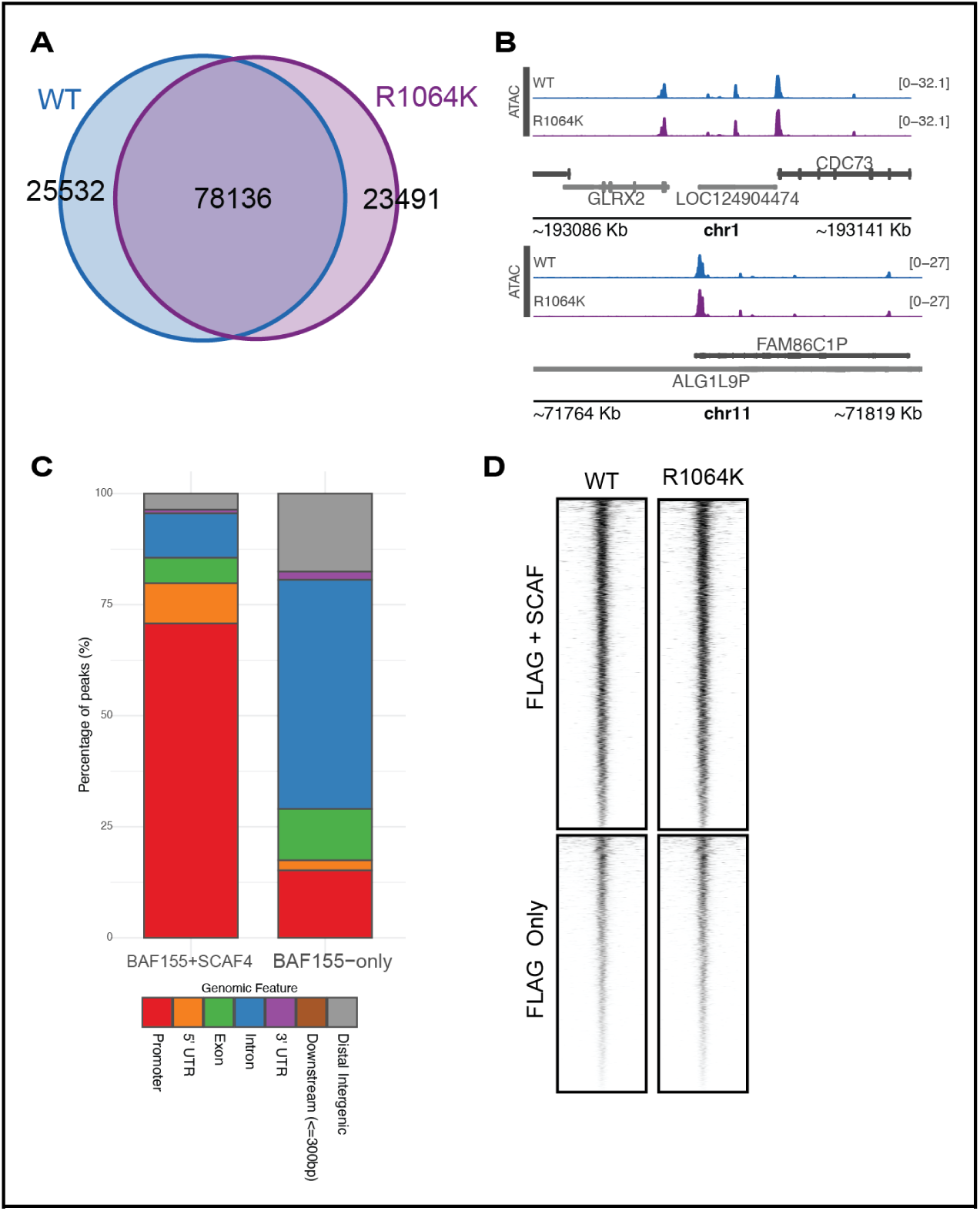
Chromatin accessibility is stable upon loss of BAF155me2a. **A.** Venn diagram of peak overlaps for HLF cell lines (n = 78136 shared, 25532 WT Only, 23491 Mutant Only) **B.** Representative genome browser tracks in BAF155-WT and BAF155-R1064K mutant cell lines. **C.** Genomic feature associated with accessible chromatin overlapping BAF155+SCAF4 or BAF155 only regions. **D.** Heatmap of accessibility signal aligned to the center point of BAF155 only or BAF155+SCAF4 peaks)

The lack of change in accessibility and genomic targeting was unexpected since BAF155 is a shared member of a chromatin remodeling complex family and forms the core heterodimer with BAF170 that forms during the initial step of complex assembly [11]. It has been previously reported that EP400/TIP60 can compensate for SWI/SNF loss in cancer cell lines [26]. We considered that a compensatory mechanism may occur when BAF155 is lost to prevent widespread transcriptional perturbation. To determine if a compensatory mechanism was at play, we performed an IP for BAF155 in WT HLF cells, BAF155-KO HLF cells, and following re-expression BAF155-FLAG HLF cell lines and immunoblotted for various SWI/SNF complex members. We observed maintained complex assembly in the absence of BAF155, most likely through compensatory expression and homodimerization of BAF170 (Supplemental Figure 3C). Notably, even SCAF4, which our data suggests interacts with BAF155 through R1064, remains associated with SWI/SNF in a BAF155-KO cell line. Together, our ATAC-seq and CUT&RUN results indicate that methylation status of BAF155 R1064K does not influence chromatin accessibility or lead to disrupted BAF155 or SCAF4 binding and that BAF155 loss may be compensated for by BAF170.

### Loss of R1064me2 leads to changes in the transcriptome and nascent transcription

We next investigated how loss of R1064 asymmetric dimethylation on BAF155 impacts gene expression since we observed an enhanced interaction between BAF155me2a and RNA processing machinery. To interrogate this, we performed mRNA-sequencing in our FLAG-tagged WT or R1064K BAF155 HLF cell lines. Differential expression analysis between the BAF155-WT and BAF155-R1064K mutant cell lines revealed 147 genes upregulated and 108 genes downregulated (padj < 0.05 & |Log2FC| > 0.6, Figure 4A, B). Without considering a fold change threshold ∼1300 genes were differentially expressed (padj < 0.05), suggesting the effects were overall moderate in magnitude.

**Figure 4.**
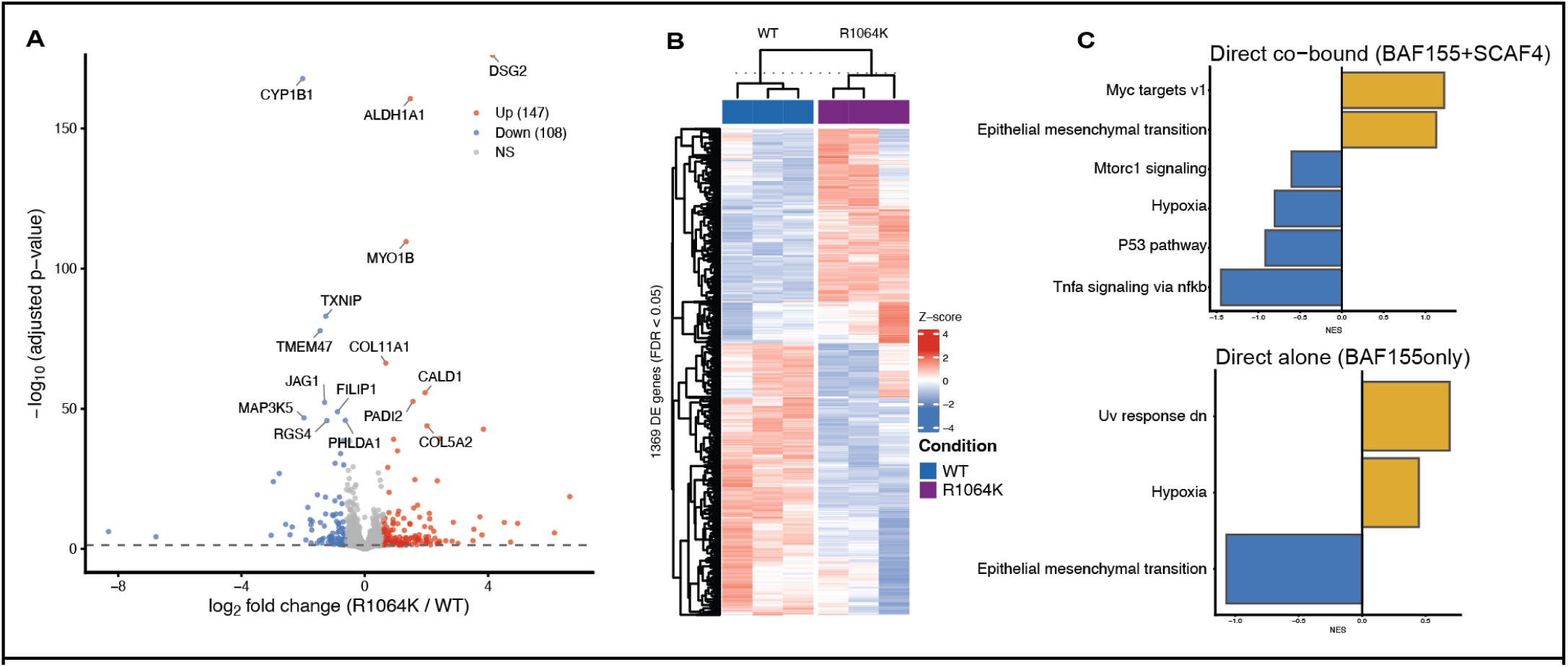
mRNA-seq reveals broad changes in transcription R1064K HLF cells. **A.** Volcano plot of top differentially expressed genes. **B.** Heatmap of differentially expressed genes (DEGs) in BAF155-FLAG WT and R1064K HLFs (padj < 0.05 & |log2FoldChange| > 0.6). **C.** Barplot of Gene Set Enrichment Analysis (GSEA) results of genes associated with BAF155+SCAF4 or BAF155 alone using the Hallmark Gene Annotation set.

We found high concordance with an orthologous RNA expression measure using 3’ end counting (Supplemental Figure 4A). GSEA analysis suggested increased expression at FLAG/BAF155 sites of Myc targets and EMT pathway genes, while genes involved in MTORC1 signaling and other signaling pathways were decreased (Figure 4C). At genes nearest to BAF155 bound without SCAF4 we observed a decrease in EMT, opposite our findings in BAF155+SCAF4 bound regions, with increased changes in hypoxia and UV response. To assess if these signaling changes were associated with altered cell growth, we measured cell proliferation in HLF BAF155-WT and BAF155-R1064K (Supplemental Figure 4B). HLF BAF155-R1064K cells grew ∼20% slower compared to BAF155-WT cells. These data suggest loss of methylation on BAF155 has moderate effects on transcription.

To determine if BAF155 methylation impacted transcription we performed transient transcriptome-sequencing (TT-seq) where nascent transcripts are transiently labelled using 4-SU and then biotinylated before isolation with streptavidin [27,28]. We used exogenous labelled yeast RNA as a spike-in control to normalize between samples. We focused analysis on genes with clear overlap between BAF155+SCAF4 or BAF155 only and removed genes with divergent, overlapping, or near multiple peaks that could not be assigned. These data revealed decreases at the 5’ end of genes on the sense strand as well as a reduction in antisense upstream transcripts (PROMPTs) at both BAF155+SCAF4 bound regions as well as BAF155-bound regions alone (Figure 5A). We quantitatively compared the upstream regions, 5’ proximal regions, the ratio between PROMPT and 5’ sense signal, as well as Serine 2 phosphorylated RNA Pol 2 (pRBP1-S2) by CUT&RUN (Figure 5B-E). These data showed widespread reduction in signal of the mutant compared to wildtype. Notably, controlling for the reduction in transcription on the sense strand, there was a larger reduction in PROMPTs at BAF155+SCAF4 bound regions that was not observed in the BAF155 only regions (Figure 5D). We expanded the signal analysis over the entire gene which showed clear reduction in nascent transcription in the 5’ end of the gene in R1064K mutant compared to WT cells that gradually normalized towards the transcript end site (TES) (Figure 5F). Using this set of genes, we analyzed whether BAF155 (FLAG), BAF155me2a, pRBP1-s2, or SCAF4 were decreased in BAF155-R1064K mutants at the TSS of these genes. Only BAF155me2a decreased, consistent with the mutation disrupting methylation (Supplemental Figure 5). Notably, we did not observe a decrease in pRBP1-S2 although the aggregate plots in Figure 5E suggested a statistically significant difference. This is due to the large sample size, but a small effect size (median Log2 Fold Change, −0.07 and −0.11 respectively). Together these data support a decrease in nascent transcription near the 5’ end of genes and decreased upstream antisense transcription in BAF155-R1064K mutants, despite normal recruitment of SCAF4 and RNA polymerase II.

**Figure 5.**
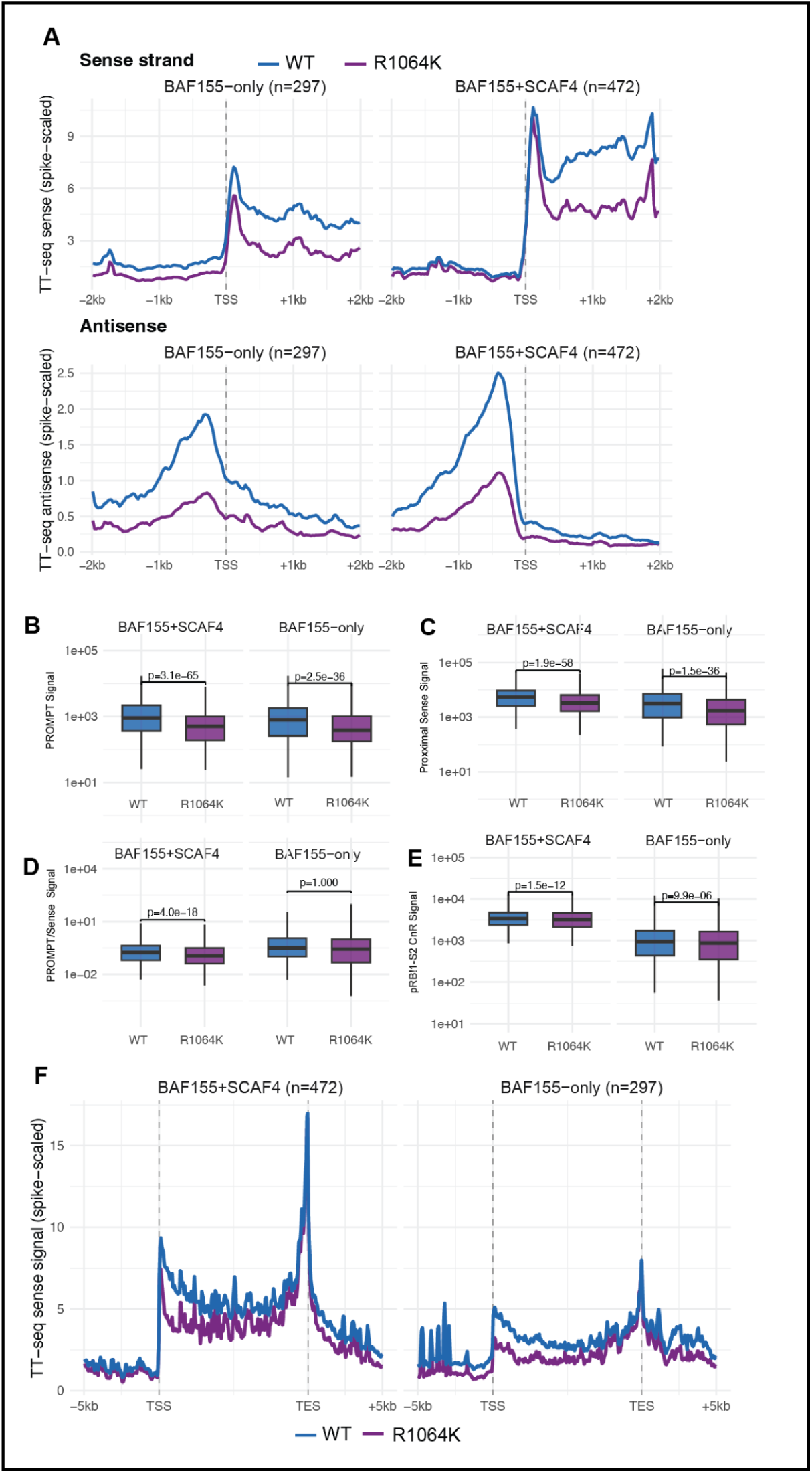
BAF155me2a loss reduces bidirectional nascent transcription. **A**. Metaplots of sense and antisense transcription at promoters of genes bound by BAF155 only or BAF155 and SCAF4. **B-D**. Quantitation of PROMPT (B), Proximal sense (C), or the ratio of PROMPT/Sense transcription (D). Quantification of serine 2 phosphorylated RBP1 at the promoter proximal region (E). **F**. Metaplots of scaled gene bodies bound by BAF155 only or BAF155 and SCAF4 depicting gradual recovery of signal at 3’ of the gene.

## Discussion

The role of PTMs on non-histone substrates and their functional implications continue to be underexplored and poorly understood. We show that asymmetric dimethylation of arginine 1064 on BAF155 influences SWI/SNF complex function. Using biochemical and multi-omics approaches, we found that modified BAF155 influences nascent transcription independently of chromatin accessibility and BAF155 binding to the genome. This result was unexpected given the well-defined function of the SWI/SNF complex in remodeling the genome through nucleosome sliding and eviction. Our genomics data instead support a model where most SWI/SNF complexes contain asymmetric dimethylated arginine and bind both promoters and distal regulatory elements.

Our data are most consistent with a defect in productive elongation. At active, BAF155-bound promoters, we observe a decrease in bidirectional nascent transcription without a corresponding decrease in RNA polymerase II loading, chromatin accessibility, or H3K27ac, suggesting that BAF155me2a functions downstream of polymerase recruitment and chromatin opening. The gradual recovery of nascent signal across the gene body, with transcription levels at the TES similar to wild type, further supports this interpretation. Polymerases that escape the promoter-proximal window appear to proceed and terminate normally. This decrease was quantitatively greater in SCAF4 bound promoters supporting a functional role for SCAF4 at these sites that is downstream of recruitment.

To identify candidate effectors of this defect, we used immunoprecipitation and mass spectrometry, which identified a large number of proteins involved in RNA processing preferentially associated with the dimethylated form of BAF155, including 3′ end cleavage and anti-terminating factors, splicing factors, and proteins that regulate the 5′ end of transcripts. We hypothesize that methylation on BAF155 bolsters these interactions and that PTMs on SWI/SNF subunits play a role in the recruitment of co-transcriptional processing machinery. We focused on the novel interactions with BAF155me2a-containing SWI/SNF complexes and SCAF4, which co-occupied promoters with BAF155. Despite co-immunoprecipitation demonstrating this interaction required BAF155 R1064, SCAF4 recruitment to chromatin was unaltered in BAF155-R1064K mutants, suggesting additional RNA processing proteins are involved with the transcriptional defects. The interplay between modified SWI/SNF and RNA processing machinery is consistent with prior work showing that SWI/SNF ATPase subunits BRG1 and BRM can influence splicing through the recruitment of splicing components, slow RNA Pol II elongation, and alter interactions with pre-mRNPs [29–31], with the established relationship between SWI/SNF complexes and RNA Pol II engagement [32], and with reports that specific subunits of SWI/SNF can both positively and negatively impact RNA polymerase pausing [33,34]. Our approaches cannot distinguish between several possibilities for how BAF155me2a impacts both sense and antisense transcripts at these promoters. However, our IP-MS data highlight several protein complexes known to act at this step, including Integrator, the cleavage and polyadenylation complex, and SCAF4 itself [24,35,36]. A shift in the balance of recruitment of these complexes, each of which can promote premature termination of early Pol II transcription, would be expected to produce the coordinated reduction of PROMPTs and 5′ sense signal that we observe.

The ability of the SWI/SNF complex to maintain a stable chromatin environment despite disruption to a core subunit highlights potential compensation between BAF155 and BAF170. Given the necessity of SWI/SNF during development and the prevalence of mutations in SWI/SNF subunits in cancer [12,13], our findings support the idea that the SWI/SNF complex has mechanisms in place to ensure that genomic regions remain open and others do not, even under prolonged perturbation. Our relatively modest effects on transcription and minimal effects on chromatin are consistent with this compensation. In BAF155-knockout cells, immunoprecipitation of BRG1 still recovers BAF170 and other core subunits (Supplemental Figure 3C), indicating that the residual complex assembles in the absence of BAF155. SCAF4 likewise co-purifies with this BAF155-less complex, providing a plausible explanation for why we do not observe a reduction in SCAF4 chromatin binding in R1064K cells: the BAF155me2a-SCAF4 contact is only one of several interfaces tethering SCAF4 to SWI/SNF, and the others are sufficient to maintain genome-wide recruitment. We interpret the partial nature of our phenotype as reflecting these multivalent interactions, which are not fully abrogated by the loss of a single PTM. Time-resolved studies will be needed to dissect the specific regulatory steps to which PTMs on SWI/SNF contribute and their role in recruiting RNA processing machinery to nascent transcription.

The number and variety of identified PTMs on non-histone substrates continue to grow and raise the need for further investigation. This study emphasizes the importance of understanding the role of PTMs on conserved chromatin remodelers and how these modifications can inform complex function. Polycomb repressive complex member EZH2 is extensively modified across cancer types [37], promoting or inhibiting metastasis through changes in protein stability depending on tissue context [38,39]. EZH2 acetylation by p300/CBP in lung adenocarcinoma is also correlated with increased invasion and metastasis through enhancer target gene repression [40]. p300 itself is auto-acetylated on its histone acetyltransferase domain to modulate enzymatic activity by displacing an autoinhibitory loop [41]. The prevalence of PTMs on broad chromatin regulators including the SWI/SNF complex suggests the significance of these modifications in guiding complex function and outcomes in both normal and disease contexts. Our data highlights regulation of RNA processing as another mode where non-histone PTMs can perform regulatory activities.

## Methods

### Key Resources Table

| Resource | Source | Catalog | Identifier |
| --- | --- | --- | --- |
| HepG2 | ATCC |  |  |
| HEK293T | Takara |  | RRID:CVCL_0045 |
| HLF |  |  | RRID:CVCL_2947 |
| lentiCrispr v2 | Gift from Feng Zhang | 52961 | RRID:Addgene_52961 |
| psPAX2 | Gift from Didier Trono | 12260 | RRID:Addgene_12260 |
| pMD2.G | Gift from Didier Trono | 12259 | RRID:Addgene_12259 |
| pFAST-BAC<br>BAF155/SMARCC1-His | Gift from Yoshihiro<br>Izumiya | 177863 | RRID:Addgene_177863 |
| cOmplete EDTA-free<br>protease inhibitor | Roche | 5056489001 |  |
| Concanavalin A Beads | Bangs Laboratories | BP531 |  |
| Protein G Magnetic<br>Beads | Cell Signaling<br>Technology | 70024 |  |
| Protein A Magnetic<br>Beads | Cell Signaling<br>Technology | 73778 |  |
| pAG-MNase | Gift from Steven Henikoff | Protein purified from plasmid in-house | RRID:Addgene_123461 |
| SilverQuest Silver Staining Kit | ThermoFisher Scientific | LC6070 |  |
| Guinea Pig anti-rabbit | Novus | NBP1-72763 | RRID:AB_11024108 |
| Goat anti-rabbit 800 | Licor | 926-32211 | RRID:AB_621843 |
| Goat anti-mouse 680 | Licor | 926-68020 | RRID:AB_10706161 |
| β-Actin (8H10D10) | Cell Signaling Technology | 3700 | RRID:AB_2242334 |
| SMARCC1/BAF155 (D7F8S) | Cell Signaling Technology | 11956 | RRID:AB_2797776 |
| Asymmetric Dimethyl-SMARCC1/B AF155 (Arg1064) (D8I3U) | Cell Signaling Technology | 94962 | RRID:AB_2800238 |
| SFRS15/SCAF4 | ThermoFisher Scientific | PA5-53780 | RRID:AB_2647188 |
| RFX5 | Sigma-Aldrich | HPA018519 | RRID:AB_1856209 |
| IgG (DA1E) | Cell Signaling Technology | 3900 | RRID:AB_1550038 |
| SMARCC2/BAF170 (D8O9V) | Cell Signaling Technology | 12760 | RRID:AB_2798017 |
| BRD9 (E9R2I) | Cell Signaling Technology | 58906 | RRID:AB_3096175 |
| SMARCE1/BAF57 (E6H5J) | Cell Signaling Technology | 33360 | RRID:AB_2799034 |
| DYKDDDDK Tag (D6W5B) (FLAG) | Cell Signaling Technology | 14793 | RRID:AB_2572291 |

### Cell Culture

All cell lines were cultured at 37°C with 5% CO_2_ in a humidified incubator. All cell lines were cultured in DMEM (Gibco) supplemented with 10% FBS and 1% Penicillin-Streptomycin (Gibco).

### Cloning

BAF155 insert was isolated from pFAST-BAC BAF155/SMARCC1-His (Addgene, 177863) [42] using PCR and cloned into a pCMV5 backbone isolated from pCMV5 BRG1-Flag (Addgene, 19143). R1064K mutation was introduced to that BAF155 insert in the pCMV5 backbone using PCR and *In Vivo* Assembly (IVA) cloning [43]. FLAG epitope was cloned onto the C-terminus of BAF155 WT and R1064K inserts in the pCMV5 constructs using IVA cloning. BAF155 WT-FLAG and R1064K-FLAG inserts were cloned into a lentiCrispr-V2 (Addgene, 52961) [44] backbone that was digested with BamHI and Xhol. Endogenous BAF155 was knocked out using the Synthego BAF155 KO kit as described [45]. Lentivirus packaging was done using OptiMEM, polyethylenimine (PEI), and a 2:2:1 ratio of psPAX2:plentiCrispr-V2:pMD2.G in HEK293T cells. Lentivirus was harvested after 72 hours. BAF155 KO cells were clonally expanded and verified using Sanger sequencing. BAF155 WT-FLAG and BAF155 R1064K-FLAG were reconstituted in BAF155 KO cells using lentiviral transduction and treated with puromycin (1 μg/mL) 72 hours after initial transduction to select for positive cells. HEK293T BAF155 WT-FLAG and BAF155 R1064K-FLAG overexpression cell lines were generated by transfecting cells with PEI, OptiMEM, and the overexpression plasmid of interest. All guide RNA and primer sequences are listed below.

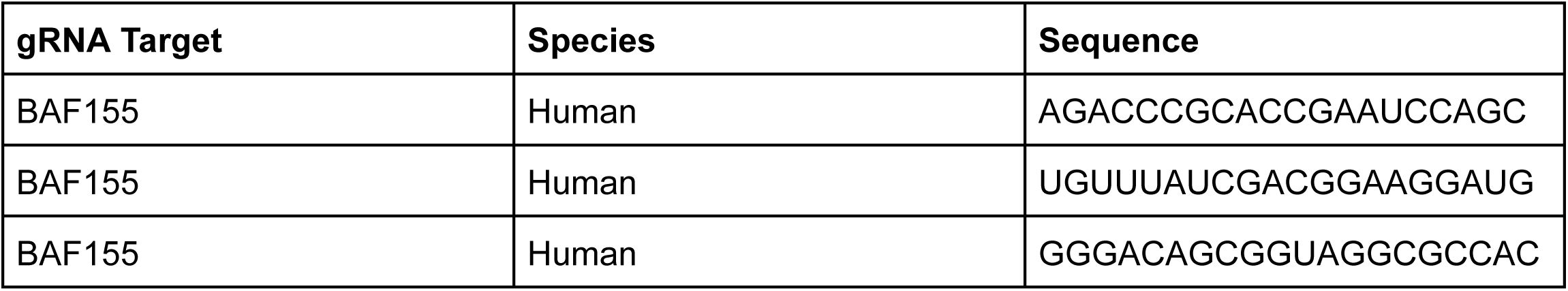

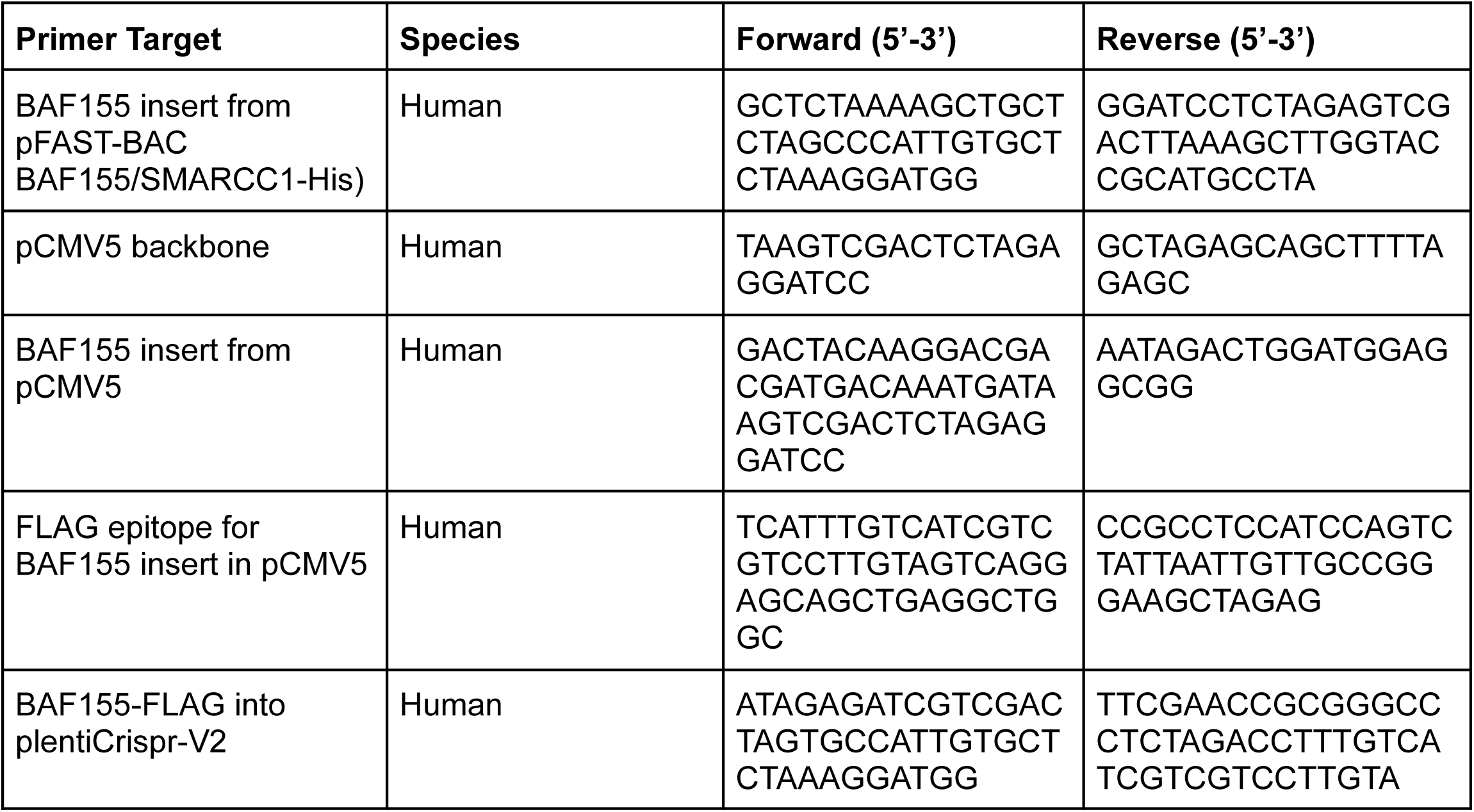
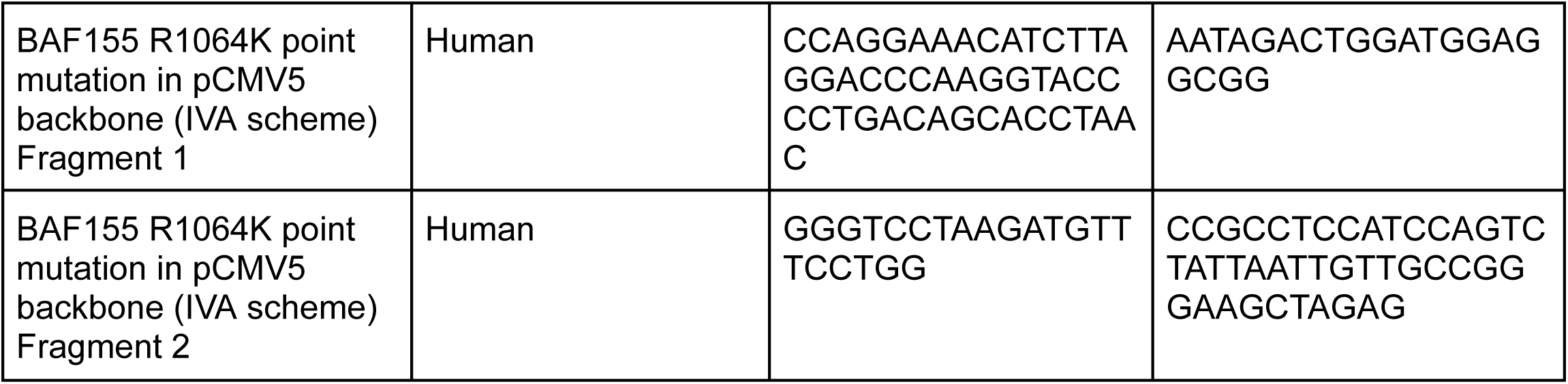

### Immunoblotting

Cells were lysed with RIPA Buffer (150 mM NaCl, 1% NP-40, 0.5% sodium deoxycholate, 1% SDS, 50 mM Tris-HCl pH 7.4, 1X PIC, 11 mM DTT, 1X universal nuclease) (∼1 million cells/30 μL). Prepped samples were loaded into a 4-15% TGX precast gel (BioRad) and separated at 250 V. Samples were transferred to a PVDF membrane (BioRad) with a Trans-Turbo Turbo machine as per the manufacturer’s manual. Membranes were blocked in a 1:1 blocking solution of Intercept blocking buffer (LI-COR) and TBS (200 mM Tris, 1500 mM NaCl) at RT for 1 hour with gentle shaking. Membranes were cut for primary antibody incubation at the manufacturer’s recommended dilution in a mix of Intercept blocking buffer, TBS, and 10% Tween 20 (1:1:0.1) overnight at 4°C with gentle shaking. The next day, membranes were removed from primary and washed 3x for 10 minutes each in TBST (TBS with 0.1% Tween 20). Membranes were then incubated in a secondary antibody mixture of Intercept blocking buffer, TBS, 10% Tween 20, and 10% SDS (1:2:0.01:0.002) at RT for one hour with light shaking and protected from light. After secondary antibody incubation, membranes were washed 2x with TBST for 10 minutes and 1x with TBS for 10 minutes. Membranes were visualized on an Odyssey CLx machine (LI-COR) and processed using ImageStudio (LI-COR).

### Immunoprecipitation

Nuclear extracts were prepared by scraping cells in 1xPBS supplemented with 0.5 mM PMSF (Sigma-Aldrich, P7626). Cells were spun at 1300 rpm for 10 minutes at 4°C and washed with 10-20 packed cell volume (PCV) of cold Buffer A (10 mM HEPES-KOH pH 7.9, 1.5 mM MgCl_2_, 10 mM KCl, 0.5 mM DTT, 0.5 mM PMSF, 1x PIC (Sigma-Aldrich, P8340)). After samples sat on ice for 10 minutes, cells were spun at 1300 rpm for 10 minutes at 4°C. Samples were homogenized with 2 PCV of cold Buffer A. Samples were pelleted for 10 minutes at 1300 rpm at 4°C. Pelleted nuclei were washed with 10 PCV of cold Buffer A and spun at 5000 rpm for 10 minutes at 4°C. Samples were moved to fresh 1.5 mL microfuge tubes and treated with 0.6 volumes of Buffer C (20 mM HEPES-KOH pH 7.9, 25% Glycerol, 420 mM KCL, 1.5 mM MgCl_2_, 0.2 mM EDTA, 0.5 mM DTT, 0.5 mM PMSF, 1x PIC). Nuclei were extracted for 30 minutes at 4°C with rotation. Lysates were clarified at 14000 rpm for 10 minutes at 4°C. The supernatant was then treated with 0.6 volumes of Buffer C and extracted for another 30 minutes. Following the second extraction, samples were brought to a final concentration of 100-150 mM KCl through the addition of 2.8 volumes of Buffer D (20 mM HEPES-KOH pH 7.9, 20% Glycerol, 0.2 mM EDTA, 0.2 mM DTT, 0.5 mM PMSF, 1X PIC). Samples were spun at 14000 rpm for 10 minutes at 4°C to clear the supernatant. The resulting pellet was snap frozen using liquid nitrogen and stored at −80°C for downstream applications.

50 μL of Protein A/G beads (Cell Signaling Technologies, 73778; 70024) were washed with wash buffer (1xPBS supplemented with 0.5% BSA (ThermoFisher, 15260037)) three times for 10 minutes each wash at 4°C. Beads were resuspended to their original volume in wash buffer. Antibodies were added to the beads at their indicated concentrations and incubated for 4 hours at 4°C with rotation. After incubation, beads were washed three times with wash buffer. The final wash was removed and beads were resuspended in nuclear extract. Samples were incubated overnight at 4°C with rotation. The next day, samples were washed on a magnet twice with BC-150 (20 mM HEPES-KOH pH 7.9, 0.15 M KCl, 10% Glycerol, 0.2 mM EDTA pH 8.0, 0.1% Tween-20, 1xPIC, 0.5 mM DTT, 0.5 mM PMSF) for 5 minutes, three times for 5 minutes with BC-300 (20 mM HEPES-KOH pH 7.9, 0.3 M KCL, 10% Glycerol, 0.2 mM EDTA pH 8.0, 0.1% Tween-20, 1xPIC, 0.5 mM DTT, 0.5 mM PMSF), and two times for 5 minutes with BC-150. Samples were moved to a new 1.5 mL microfuge tube for the final 5 minute was with BC-FINAL (20 mM HEPES-KOH pH 7.9, 0.06 M KCl, 10% Glycerol, 1xPIC, 0.5 mM DTT, 0.5 mM PMSF). Samples were eluted using 1xLaemmli buffer (17 mM DTT, 50 mM Tris-Cl pH 6.8, 2% SDS, 0.1% bromophenol blue) and shaken at 1500 rpm for 5 minutes at 37°C in a thermomixer. Samples were placed on a magnet and the supernatant was moved to a new 1.5 mL microfuge tube. Lysates were incubated at 100°C for 10 minutes. Samples were stored at −20°C for downstream applications.

### Silver Stain

Silver staining was performed using the SilverQuest Silver Staining Kit (ThermoFisher, LC6070). Briefly, cells were lysed with RIPA Buffer (∼1 million cells/30 μL). Prepped samples were loaded into a 4-15% TGX precast gel (BioRad) and separated. Following electrophoresis, the gel was removed from the cassette and placed in a clean tray. The gel was briefly washed with ultra-pure water. The gel was incubated with 100 mL of Fixative (40% ethanol, 10% acetic acid, made with ultra-pure water) for 20 minutes at room temperature with gentle rotation. The gel was then incubated with 30% ethanol for 10 minutes at room temperature, followed by a 10 minute incubation with 100 mL Sensitizing Solution (30% ethanol, 10 % Sensitizer, ultra-pure water). The gel was incubated with 30% ethanol for 10 minutes at room temperature and then washed with 100 mL ultra-pure water for 10 minutes. The gel was stained by incubation with 100 mL Staining Solution (1% Stainer, ultra-pure water) for 15 minutes at room temperature. The gel was then briefly washed with ultra-pure water for 20-60 seconds and incubated with 100 mL Developing Solution (10% Developer, 1 drop Developer enhancer, ultra-pure water) for 4-8 minutes to visualize the signal. Once the desired intensity was met, 10 mL of Stopper was added directly to the solution and shaken gently for 10 minutes at room temperature. A final wash in 100 mL ultra-pure water for 10 minutes at room temperature was performed before imaging the gel.

### mRNA-seq

Total RNA was isolated from cultured cells using Monarch Total RNA Miniprep Kit (New England Biolabs, T2010S) following the manufacturer’s instructions. RNA concentration and purity were determined using Agilent TapeStation systems (Agilent, 5067-5579). Samples with RNA integrity number (RIN) ≥ 8 were used for library preparation. Samples were prepared for strand specific RNA-seq and sequenced on a Novaseq machine at 2×150bp by Admera.

### CUT&RUN

CUT&RUN protocol was adapted from Epicypher and [46]. Briefly, cells were detached from plates and counted for a total of ∼500,000 cells per antibody per cell line. Cells were pelleted at RT for 3 minutes at 300 g. Cells were washed 2x with Wash Buffer (20mM HEPES pH 7.6, 150mM NaCl, 0.5mM Spermidine, protease inhibitor (1 tablet/30mL Roche)). 11 μL of Concanavalin beads (Bangs Laboratories) per sample were 2x washed with cold Bead Activation Buffer (20mM HEPES ph7.9, 10mM KCl, 1mM CaCl2, and 1mM MnCl2). Cells were then added to beads and incubated with rotation for 5 minutes at RT. Samples were placed on a magnet to remove the supernatant and resuspended in 50 μL Wash Buffer with 0.05% Digitonin and 2mM EDTA. Corresponding antibodies were added to the samples and were incubated overnight at 4℃ with rotation. The following day, beads were washed on a magnet 3x with Digitonin-Wash Buffer (Wash Buffer, 0.05% Digitonin).

Beads were resuspended in 50 μL Digitonin-Wash Buffer with guinea pig anti-rabbit secondary antibody (1:100, Novus, NBP1-72763) and incubated at 4℃ for 1 hour with rotation. Following secondary antibody binding, beads were washed 4x with cold Digitonin-Wash buffer on a magnet and then resuspended in 50 μL pAG Mnase solution (∼700 ng/ul pAG-Mnase, Digitonin-Wash Buffer) (purified in-house [47]). Beads were incubated with pAG Mnase solution for 1 hour at 4℃ with rotation. Beads were washed 4x with cold Digitonin-Wash Buffer on a magnet and then resuspended in 50 μL cold Digitonin-Wash Buffer. 1 μL of 100 mM CaCl_2_ was added to the samples and incubated in a cold metal block in ice at 4℃ for 30 minutes. 50 μL of Stop Buffer (340mM NaCl, 20mM EDTA, 4mM EGTA, 0.05µg/µL RNAseA (ThermoFisher, EN0531), 0.1% Triton X-100) was added to the samples and then incubated at 37℃ for 30 minutes in a thermocycler. DNA was purified from the samples using the Zymo ChIP Clean and Concentrator kit (Genesee Scientific, 11-379C) following the manufacturer’s protocol. Libraries were prepared using the KAPA HyperPrep DNA kit with modifications. End-repair/A-tailing was performed in 25 μL volume reactions and performed on a thermocycler with the following protocol: 12℃ for 15 minutes, 37℃ for 15 minutes, 58℃ for 45 minutes, hold at 12℃. For adapter ligation, half volumes were used with 5 μL of 750 nM Kapa Unique Dual-Indexed Adapters (KAPA, 8861919702). Samples were incubated for 1 hour at RT and cleaned with a 1.1X KAPA Pure Bead clean up 2x. A library amplification was performed with KAPA HiFi HotStart ReadyMix with the following program: 98℃ for 45 seconds, 98℃ for 15 seconds, 60℃ for 10 seconds, Cycle to Step 2 13x, 71℃ for 60 seconds, hold at 4℃. A final KAPA Pure Bead 1X clean up was performed on the amplified libraries before being quantified and pooled on TapeStation 4200 and Qubit. Samples were sequenced using Illumina sequencing on a Novaseq X 2×150bp.

### ATAC-seq

ATAC-seq was performed using the Omni-ATAC protocol [48]. Briefly, 50,000 viable cells per sample were pelleted at 500 × g for 5 minutes at 4°C. Cell pellets were resuspended in 50 μL of cold ATAC-Resuspension Buffer (RSB) containing 0.1% IGEPAL CA-630, 0.1% Tween-20, and 0.01% digitonin, then pipetted up and down three times and incubated on ice for 3 minutes. Lysis buffer was washed out with 1 mL of cold ATAC-RSB containing 0.1% Tween-20 (without IPEGAL or digitonin), and nuclei were pelleted at 500 xg for 10 minutes at 4°C. Nuclei were resuspended in 50 μL of transposition mix containing 25 μL of 2x TD buffer (Diagenode), 0.05% digitonin, 0.1% Tween-20, and 200 nM Tn5 transposase (custom-loaded, diluted from a 37.5 μM stock). Transposition was carried out at 37°C for 30 minutes in a thermomixer at 1000 RPM. Reactions were cleaned up using the Zymo DNA Clean & Concentrator-5 kit (D4014, Zymo Research) and eluted in 21 μL Elution Buffer.

Libraries were pre-amplified using NEBNext High-Fidelity 2x PCR Master Mix (NEB, M0530) with indexed primers under the following conditions: 72°C for 5 minutes, 98°C for 30 seconds, followed by 5 cycles of 98°C for 10 seconds, 63°C for 30 seconds, and 72°C for 1 minute. To determine the optimal number of additional cycles, 10% of the pre-amplified material was subjected to a qPCR side reaction following the method described by [49]. Final amplification was performed accordingly, and libraries were purified using a two-sided bead cleanup (KAPA Pure Beads, Roche) with 0.6x and 1.2x ratios to remove large fragments and unincorporated adapters. Library concentration and quality were assessed using the Agilent TapeStation with High Sensitivity D1000 ScreenTape. Libraries were pooled and sequenced on an Illumina NextSeq 1000 platform with 2×50 bp paired-end reads.

### mRNA-seq analysis

mRNA-seq FASTQ files were processed using our Nextflow RNA-seq pipeline (https://github.com/raab-lab/rnaseq). This pipeline produces transcript count matrices using Salmon (Version 1.10.0) [50]. Transcripts were imported and merged in R using the tximeta package [51].

DESeq2 was used to perform differential expression analysis in R (Version 4.3.1, unless stated otherwise) [52]. Log2FoldChange values were shrunk using lfcShrink (type = “apeglm”) [53] to reduce noise. All downstream analysis was performed in R. Gene Set Enrichment Analysis (GSEA) was performed using clusterProfiler (Version = 4.8.3) with gene annotations extracted from msigdbr (Version = 10.0.2) [54].

### TT-seq

TT-seq was performed as previously described (Gregersen et al., 2020). In short, cells were seeded at ∼80% confluency in 3, 15 cm dishes 24 hours prior to labeling. Cells were pulse-labeled with 1 mM of 4-thiouridine (4SU) for 15 minutes with labeling being quenched with the removal of the media and the addition of 2 mL of TRIzol (ThermoFisher, 15596026). RNA was extracted using phenol/chloroform and quantified on a TapeStation 4200 (Agilent, G2991BA). Incorporation of 4SU in extracted RNA was validated using dot blot and imaged with an Odyssey CLx machine (LI-COR).

Labeled RNA was fragmented using 1 M NaOH and isolated using Micro Bio-Gel spin columns (BioRad, 7326250). Fragmented RNA was then biotinylated with biotin buffer and 0.1 mg/mL MTSEA biotin-XX linker (Biotium, BT90066) for 30 minutes. RNA was harvested using phenol/chloroform/isoamyl alcohol (25:24:1 (vol/vol/vol)) followed by precipitation with 5 M NaCl and isopropanol. Biotinylated RNA was extracted from the unlabeled RNA by incubation with μMACS streptavidin MicroBeads (Miltenyi, 130-074-101) and subsequent pull-down with μColumns (Miltenyi, 130-042-701) and warmed pull down wash buffer (100 mM Tris-HCl, pH 7.4, 10 mM EDTA, 1 M NaCl, 0.1% (vol/vol) Tween 20). After pull-down, 4SU-RNA was cleaned and concentrated using the RNeasy MinElute Cleanup Kit (Qiagen, 74204). Size and quality of the isolated RNA was checked on a TapeStation 4200. Library prep was performed using the Hyperprep Kit (KAPA, 08098093702) for degraded RNA (1 minute fragmentation step) with Kapa Dual Index Adapters (KAPA, 08278555702). Size and concentration of libraries were verified using TapeStation 4200 and Qubit (Invitrogen, Q32857). Libraries were pooled and sequenced at 2×150 bp on a Novaseq X Plus 10B.

### TT-seq Analysis

TT-seq data were aligned to human genome version “hg38” and gencode v32 using STAR with the following parameters: --runMode alignReads --outSamunmapped None --twoPassMode --none. A second pass aligning reads to the yeast genome using bowtie2 and the sacCer3 UCSC genome index was performed to derived normalization factors as the sum of properly mapped reads to yeast genome from ‘samtools flagstat’ with parameters --very-sensitive --no-unal. Bigwig files were computed using deeptool bamCoverage with --scaleFactor 1/[(# of correct pairs of yeast reads)/median(# correct pairs of yeast reads)] --filterStrand [reverse|forward] --binSize 10. Metaplots and coverage over regions of interest were calculated in R.

### CUT&RUN analysis

CUT&RUN data was processed using our Nextflow CUT&RUN pipeline (github.com/raab-lab/cut-n-run Version 4.0). This pipeline performs alignment, filtering, peak calling, sorting, and produces bigwig files for peak visualization. Reads were aligned using Bowtie2 –very-sensitive-local -X 800 parameters to hg38 and trimmed using TrimGalore. Samples are then sorted and indexed using Samtools. Peak calling was performed using MACS2 (--call-summits qvalue=0.05). Duplicates were removed using Picard and peak normalization for each antibody was calculated using CSAW then applied to each track. Coverage tracks were produced using Deeptools (version 3.5.4). Final coverage tracks were produced by averaging bigwig signals across replicates.

All outlined steps were performed within the Raab Lab Nextflow CUT&RUN pipeline (raab-lab/cut-n-run). All downstream analysis was performed using R (Version 4.3.1, unless stated otherwise). Consensus peaks for each antibody across replicates was calculated using the rmspc package in R [55]. Peak annotation to the nearest gene and genomic feature was performed using the ChIPSeeker [56] and ChIPpeakAnno [56,57] packages. CUT&RUN signal heatmaps and metaplots were generated using computeMatrix and plotHeatmap functions from Deeptools. Differential binding analysis on consensus peaks was performed using the csaw package [58,59]. Transcription factor motif analysis was performed using HOMER [60].

### ATAC-seq analysis

Fastq files were processed using our Nextflow pipeline (github.com/raab-lab/cut-n-run –atac). This pipeline is similar to our CUT&RUN approach, with some ATAC-seq specific changes denoted by the –atac flag. Briefly, reads are aligned with Bowtie2, mitochondrial reads are filtered, duplicates are marked with PICARD, and normalization factors are calculated using CSAW ‘efficiency’ approach on a consensus set of peaks from all replicates/conditions defined using MSPC. For differential accessibility, reads overlapping consensus peaks were counted using the CSAW function regionCounts and then used as input for DESeq2 to perform differential testing and lfcShrink(type = ‘apeglm’), but this yielded no statistically significant changes. Metaplots of ATAC-seq signal were calculated in R on coverage matrices produced using deeptools computeMatrix.

### Cell Proliferation Assay

Cells were seeded at ∼5,000 cells per well in triplicate in a 96 well plate and incubated for 4 days. On day 4, cell growth was measured using CellTiter-Glo® (CTG) luminescent cell viability assay (Promega) with a 1:1 ratio of CTG reagent for 10 minutes at RT with gentle shaking. Signal from samples was then measured using a fluorescence assay on Cytation 5 machine (BioTek). Growth was plotted using R (Version 4.3.1) with relative fluorescence ± standard deviation.

## Data Availability

All sequencing data will be deposited at GEO upon manuscript submission for review. Tables of processed data are available in supplemental files. Code will be deposited at github.com/raab-lab/sokolowski_baf155_2026. Any requests for additional materials or data will be fulfilled by the corresponding author

## AI Usage Statement

Claude Opus 4.6/4.7 and Sonnet 4.5/4.6 were used in the generation of some code following example scripts and drafts written initially by lab members. Some text editing was aided through the use of these models. The authors are responsible for all final versions.

## Author Contributions

Mallory Sokolowski: Formal analysis, Investigation, Data curation, Visualization, Methodology, Validation, Writing—original draft, review & editing. Deena Scoville: Investigation, Formal analysis, Methodology. Peyton Kulers: Data curation, Software, Investigation; Jesse Raab: Conceptualization, Investigation, Formal analysis, Data curation, Methodology, Visualization, Software, Supervision, Writing—original draft, review & editing.

## Supplementary Figures

**Supplemental Figure 1.**
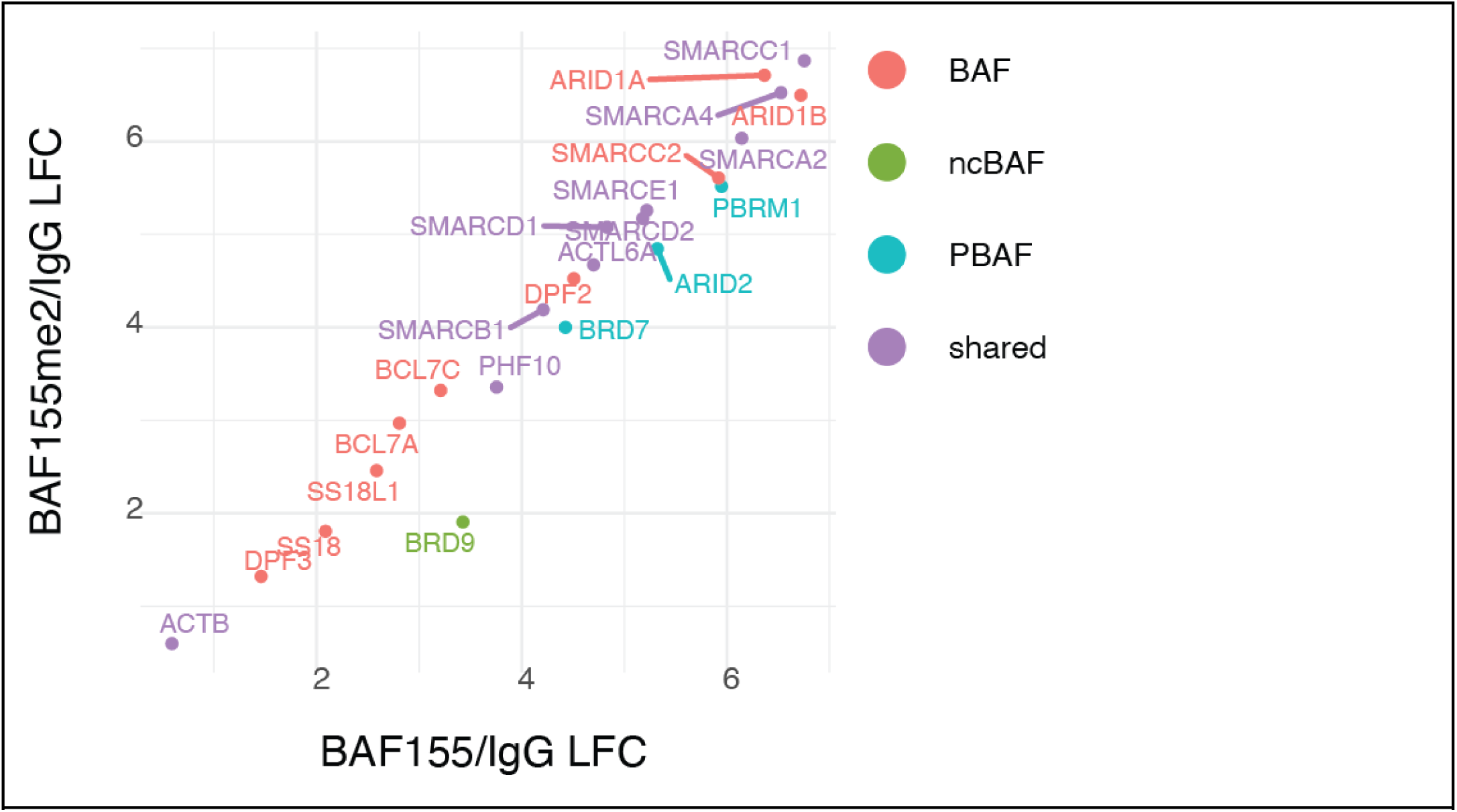
BAF155me2a does not impact types of SWI/SNF complexes. Log2 Fold Change ratios of IP relative to non-specific IgG IP for total BAF155 antibody (x-axis) compared to BAF155me2a antibody (y-axis). Proteins are colored by SWI/SNF subcomplex family assignment.

**Supplemental Figure 2.**
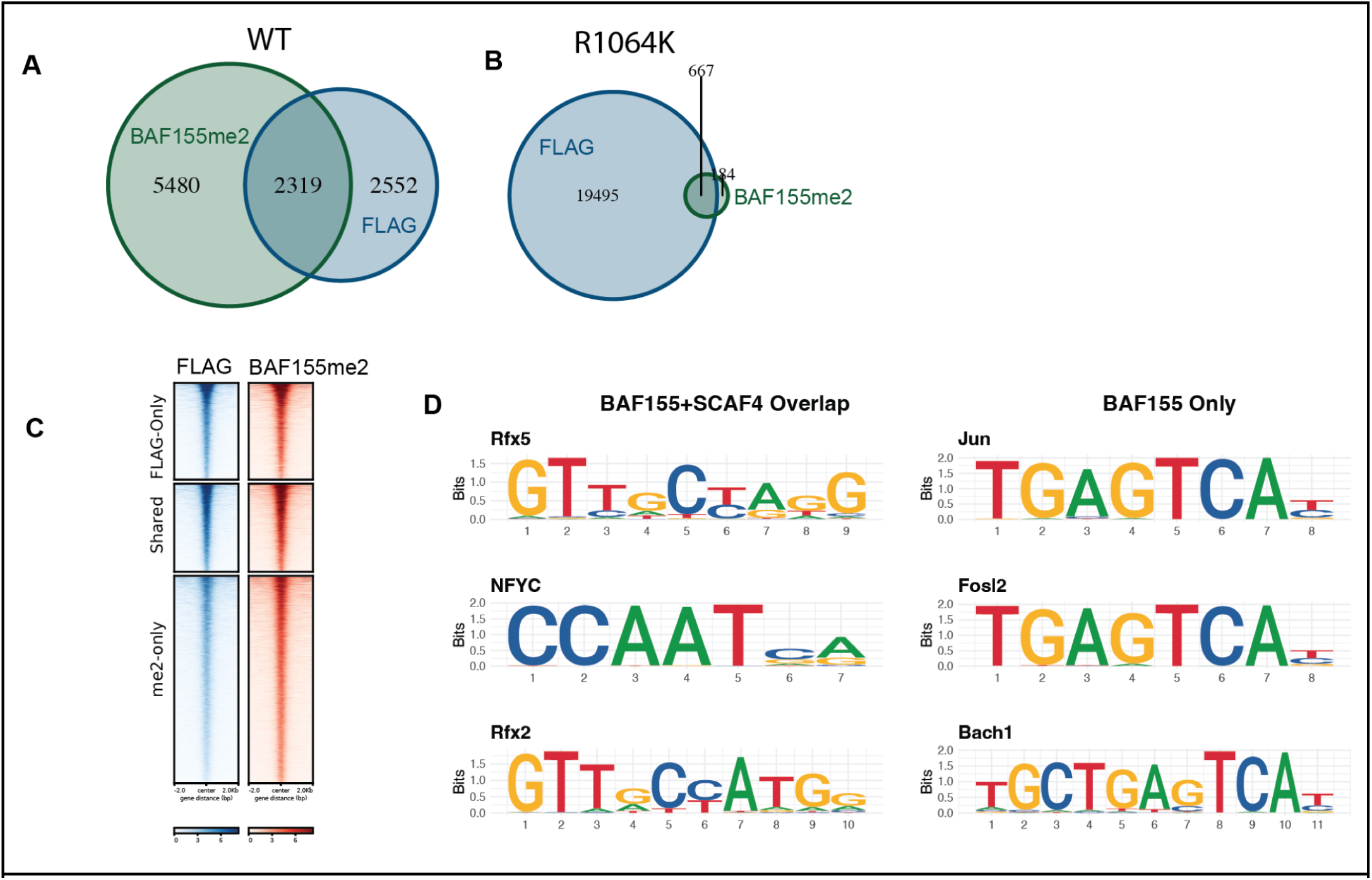
BAF155me2 and Total BAF155 overlap throughout the genome. **A-B**. Venn diagram of peak calls in (A) WT cells or BAF155-R1064K mutants (B) for BAF155me2a and BAF155-FLAG. **C**. Heatmap of signal intensity for BAF155-FLAG or BAF155me2a at venn diagram regions in A. **D**. Example motifs enriched in either BAF155+SCAF4 or BAF155 only sites.

**Supplemental Figure 3.**
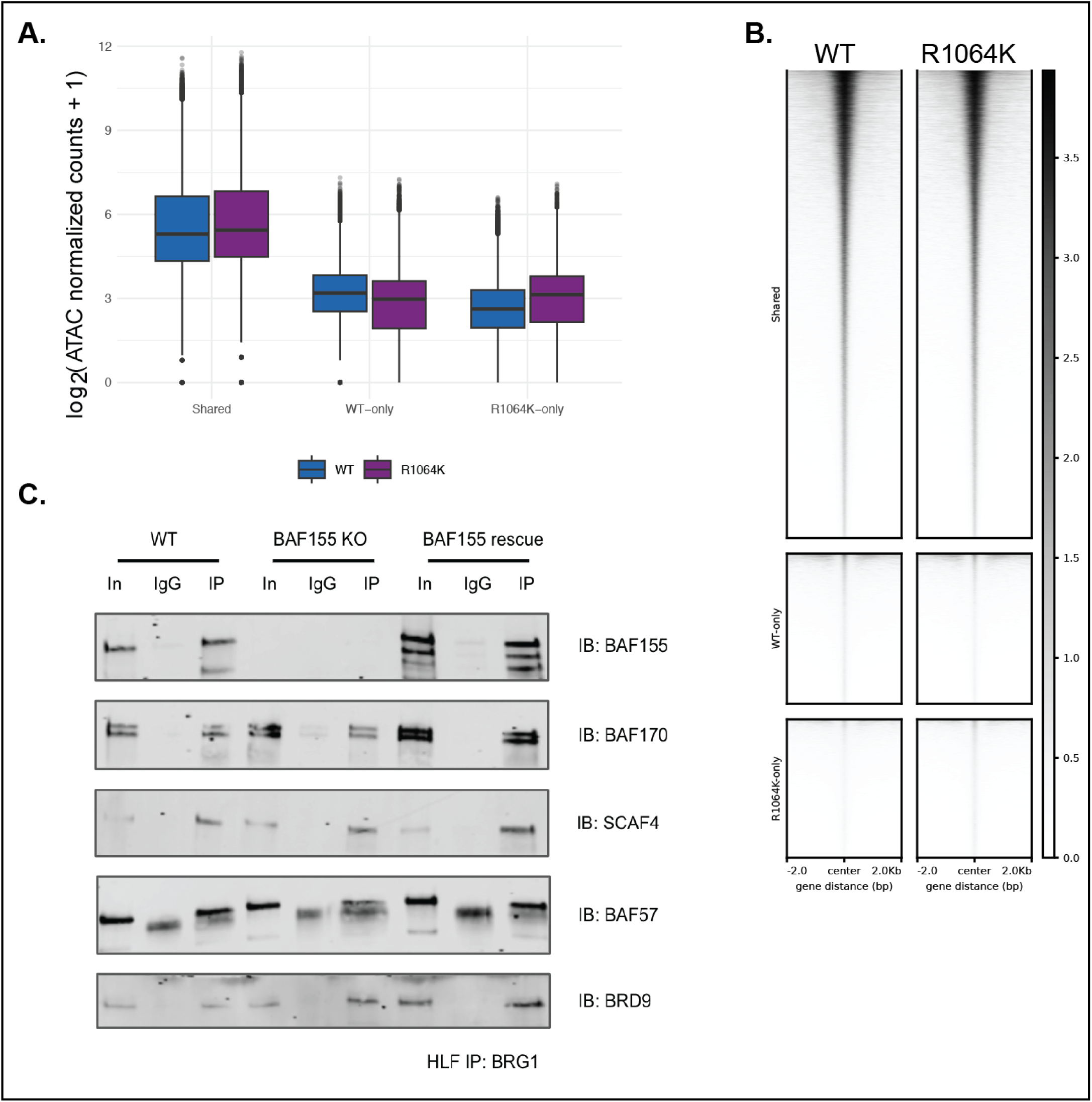
Chromatin accessibility does not change in BAF155 R1064K mutant. **A**. Quantitation of ATAC-seq signal at ‘Shared’, ‘WT-Only’, or ‘R1064K-Only’ accessible regions. **B**. Heatmap of ATAC-seq signal at three categories of open chromatin regions from A. **C**. Immunoprecipitations in HLF cell lines using BRG1 antibody followed by immunoblotting for SWI/SNF subunits and SCAF4 depicting interactions are maintained in the absence of BAF155.

**Supplemental Figure 4.**
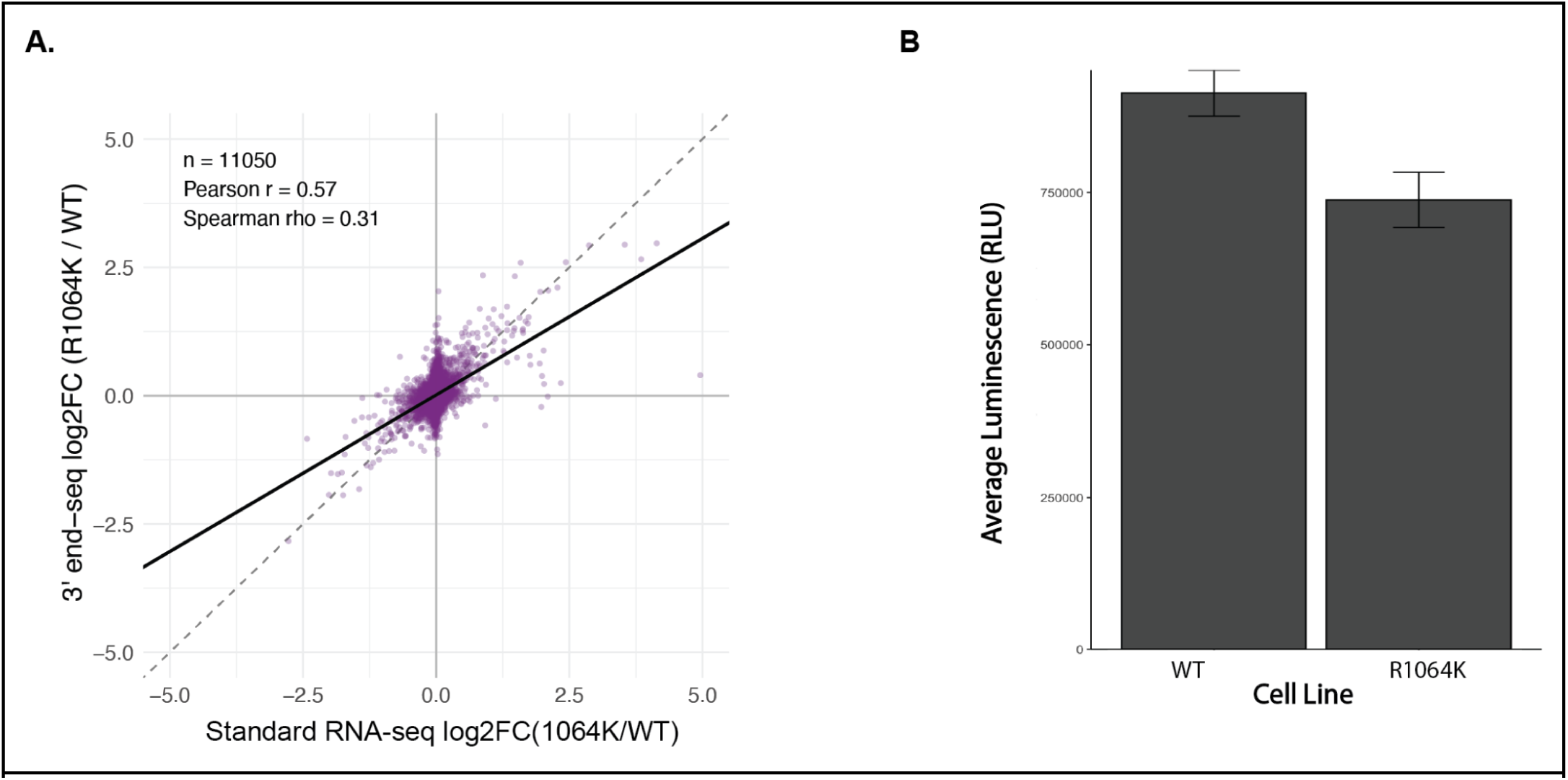
Expression and proliferation analysis. **A**. Correlation plot of standard strand-specific RNA seq (x-axis) compared to 3’ end counting RNA-seq (Plasmidsaurus, y-axis). **B**. Cell growth assay using Cell Titer glow comparing WT and R1064K HLF cell lines.

**Supplemental Figure 5.**
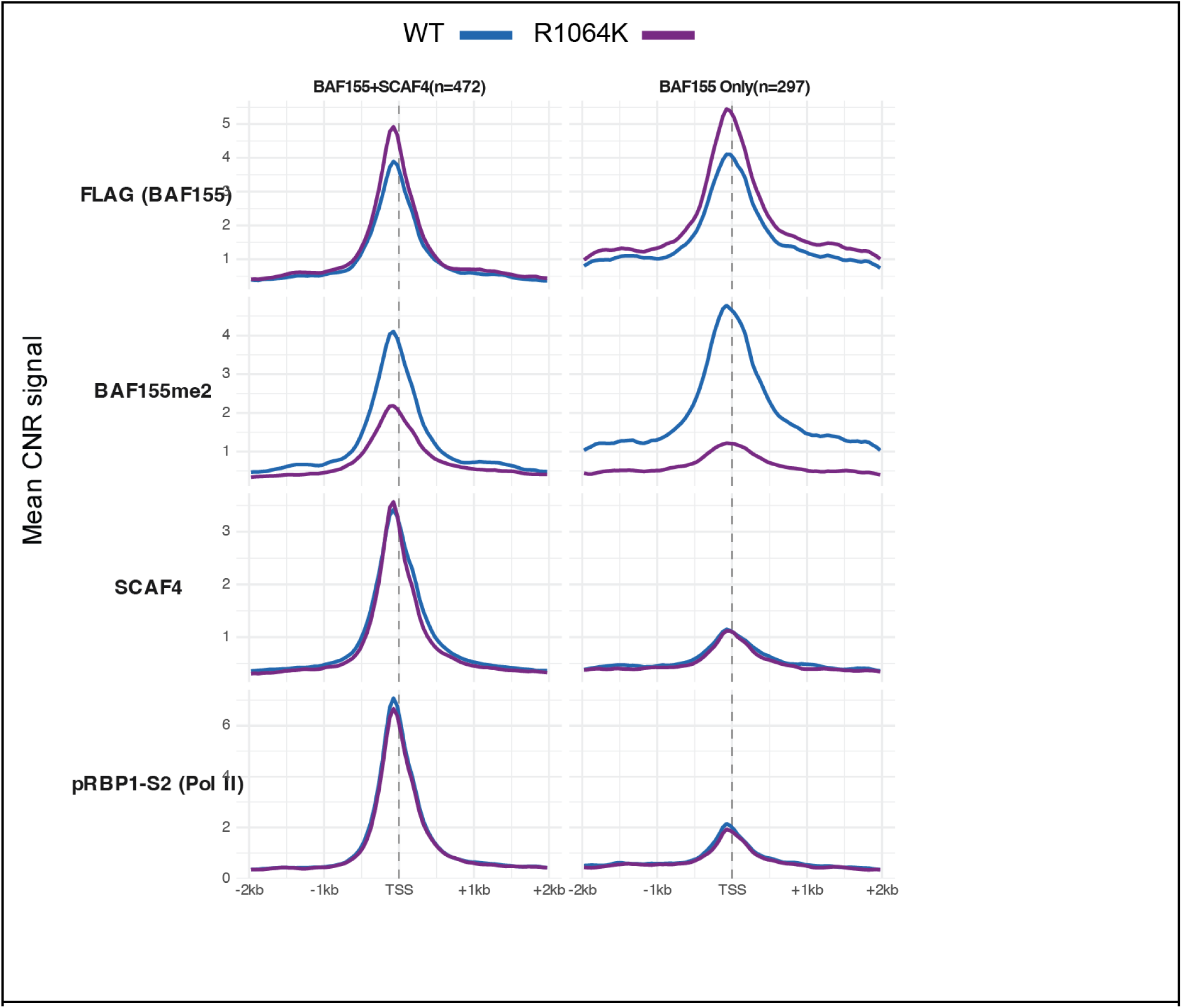
Chromatin regulator binding is unchanged at genes with decreased nascent transcription. Metaplots for BAF155-FLAG, BAF155me2a, SCAF4, or serine 2 phosphorylated RBP1 at 5’ TSS of promoters. Only BAF155me2 signal decreases in the mutant.

## References

1. Clapier CR, Cairns BR. The biology of chromatin remodeling complexes. Annu Rev Biochem. 2009;78: 273–304. doi:10.1146/annurev.biochem.77.062706.153223

2. Ho L, Crabtree GR. Chromatin remodelling during development. Nature. 2010;463: 474–484. doi:10.1038/nature08911

3. Ahmad K, Brahma S, Henikoff S. Epigenetic pioneering by SWI/SNF family remodelers. Mol Cell. 2024;84: 194–201. doi:10.1016/j.molcel.2023.10.045

4. Hodges C, Kirkland JG, Crabtree GR. The Many Roles of BAF (mSWI/SNF) and PBAF Complexes in Cancer. Cold Spring Harb Perspect Med. 2016;6. doi:10.1101/cshperspect.a026930

5. Clapier CR, Iwasa J, Cairns BR, Peterson CL. Mechanisms of action and regulation of ATP-dependent chromatin-remodelling complexes. Nat Rev Mol Cell Biol. 2017;18: 407–422. doi:10.1038/nrm.2017.26

6. Narlikar GJ, Sundaramoorthy R, Owen-Hughes T. Mechanisms and functions of ATP-dependent chromatin-remodeling enzymes. Cell. 2013;154: 490–503. doi:10.1016/j.cell.2013.07.011

7. Raab JR, Resnick S, Magnuson T. Genome-Wide Transcriptional Regulation Mediated by Biochemically Distinct SWI/SNF Complexes. PLoS Genet. 2015;11: e1005748. doi:10.1371/journal.pgen.1005748

8. Wang W, Côté J, Xue Y, Zhou S, Khavari PA, Biggar SR, et al. Purification and biochemical heterogeneity of the mammalian SWI-SNF complex. EMBO J. 1996;15: 5370–5382. Available: https://www.ncbi.nlm.nih.gov/pubmed/8895581

9. Alpsoy A, Dykhuizen EC. Glioma tumor suppressor candidate region gene 1 (GLTSCR1) and its paralog GLTSCR1-like form SWI/SNF chromatin remodeling subcomplexes. J Biol Chem. 2018;293: 3892–3903. doi:10.1074/jbc.RA117.001065

10. Gatchalian J, Malik S, Ho J, Lee D-S, Kelso TWR, Shokhirev MN, et al. A non-canonical BRD9-containing BAF chromatin remodeling complex regulates naive pluripotency in mouse embryonic stem cells. Nat Commun. 2018;9: 5139. doi:10.1038/s41467-018-07528-9

11. Mashtalir N, D’Avino AR, Michel BC, Luo J, Pan J, Otto JE, et al. Modular Organization and Assembly of SWI/SNF Family Chromatin Remodeling Complexes. Cell. 2018;175: 1272–1288.e20. doi:10.1016/j.cell.2018.09.032

12. Kadoch C, Hargreaves DC, Hodges C, Elias L, Ho L, Ranish J, et al. Proteomic and bioinformatic analysis of mammalian SWI/SNF complexes identifies extensive roles in human malignancy. Nat Genet. 2013;45: 592–601. doi:10.1038/ng.2628

13. Bultman S, Gebuhr T, Yee D, La Mantia C, Nicholson J, Gilliam A, et al. A Brg1 null mutation in the mouse reveals functional differences among mammalian SWI/SNF complexes. Mol Cell. 2000;6: 1287–1295. doi:10.1016/s1097-2765(00)00127-1

14. Kim JK, Huh SO, Choi H, Lee KS, Shin D, Lee C, et al. Srg3, a mouse homolog of yeast SWI3, is essential for early embryogenesis and involved in brain development. Mol Cell Biol. 2001;21: 7787–7795. doi:10.1128/MCB.21.22.7787-7795.2001

15. Gao X, Tate P, Hu P, Tjian R, Skarnes WC, Wang Z. ES cell pluripotency and germ-layer formation require the SWI/SNF chromatin remodeling component BAF250a. Proc Natl Acad Sci U S A. 2008;105: 6656–6661. doi:10.1073/pnas.0801802105

16. de la Serna IL, Ohkawa Y, Imbalzano AN. Chromatin remodelling in mammalian differentiation: lessons from ATP-dependent remodellers. Nat Rev Genet. 2006;7: 461–473. doi:10.1038/nrg1882

17. Ho PJ, Lloyd SM, Bao X. Unwinding chromatin at the right places: how BAF is targeted to specific genomic locations during development. Development. 2019;146. doi:10.1242/dev.178780

18. Larsen SC, Sylvestersen KB, Mund A, Lyon D, Mullari M, Madsen MV, et al. Proteome-wide analysis of arginine monomethylation reveals widespread occurrence in human cells. Sci Signal. 2016;9: rs9. doi:10.1126/scisignal.aaf7329

19. Padilla-Benavides T, Haokip DT, Yoon Y, Reyes-Gutierrez P, Rivera-Pérez JA, Imbalzano AN. CK2-Dependent Phosphorylation of the Brg1 Chromatin Remodeling Enzyme Occurs during Mitosis. Int J Mol Sci. 2020;21. doi:10.3390/ijms21030923

20. Guo P, Hoang N, Sanchez J, Zhang EH, Rajawasam K, Trinidad K, et al. The assembly of mammalian SWI/SNF chromatin remodeling complexes is regulated by lysine-methylation dependent proteolysis. Nat Commun. 2022;13: 6696. doi:10.1038/s41467-022-34348-9

21. Wang L, Zhao Z, Meyer MB, Saha S, Yu M, Guo A, et al. CARM1 methylates chromatin remodeling factor BAF155 to enhance tumor progression and metastasis. Cancer Cell. 2014;25: 21–36. doi:10.1016/j.ccr.2013.12.007

22. Kim E-J, Liu P, Zhang S, Donahue K, Wang Y, Schehr JL, et al. BAF155 methylation drives metastasis by hijacking super-enhancers and subverting anti-tumor immunity. Nucleic Acids Res. 2021;49: 12211–12233. doi:10.1093/nar/gkab1122

23. Bajpai R, Chen DA, Rada-Iglesias A, Zhang J, Xiong Y, Helms J, et al. CHD7 cooperates with PBAF to control multipotent neural crest formation. Nature. 2010;463: 958–962. doi:10.1038/nature08733

24. Gregersen LH, Mitter R, Ugalde AP, Nojima T, Proudfoot NJ, Agami R, et al. SCAF4 and SCAF8, mRNA Anti-Terminator Proteins. Cell. 2019;177: 1797–1813.e18. doi:10.1016/j.cell.2019.04.038

25. Skene PJ, Henikoff S. An efficient targeted nuclease strategy for high-resolution mapping of DNA binding sites. Elife. 2017;6. doi:10.7554/eLife.21856

26. Martin BJE, Ablondi EF, Goglia C, Mimoso CA, Espinel-Cabrera PR, Adelman K. Global identification of SWI/SNF targets reveals compensation by EP400. Cell. 2023;186: 5290–5307.e26. doi:10.1016/j.cell.2023.10.006

27. Schwalb B, Michel M, Zacher B, Frühauf K, Demel C, Tresch A, et al. TT-seq maps the human transient transcriptome. Science. 2016;352: 1225–1228. doi:10.1126/science.aad9841

28. Gregersen LH, Mitter R, Svejstrup JQ. Using TT-seq for profiling nascent transcription and measuring transcript elongation. Nat Protoc. 2020;15: 604–627. doi:10.1038/s41596-019-0262-3

29. Batsché E, Yaniv M, Muchardt C. The human SWI/SNF subunit Brm is a regulator of alternative splicing. Nat Struct Mol Biol. 2006;13: 22–29. doi:10.1038/nsmb1030

30. Zapater AG, Mackowiak SD, Guo Y, Jordan-Pla A, Friedländer MR, Visa N, et al. The SWI/SNF subunits BRG1 affects alternative splicing by changing RNA binding factor interactions with RNA. bioRxiv. 2019. p. 858852. doi:10.1101/858852

31. Tyagi A, Ryme J, Brodin D, Ostlund Farrants AK, Visa N. SWI/SNF associates with nascent pre-mRNPs and regulates alternative pre-mRNA processing. PLoS Genet. 2009;5: e1000470. doi:10.1371/journal.pgen.1000470

32. Brahma S, Henikoff S. The BAF chromatin remodeler synergizes with RNA polymerase II and transcription factors to evict nucleosomes. Nat Genet. 2024;56: 100–111. doi:10.1038/s41588-023-01603-8

33. Trizzino M, Barbieri E, Petracovici A, Wu S, Welsh SA, Owens TA, et al. The Tumor Suppressor ARID1A Controls Global Transcription via Pausing of RNA Polymerase II. Cell Rep. 2018;23: 3933–3945. doi:10.1016/j.celrep.2018.05.097

34. Zhu X, Fu Z, Aceto G, St-Germain J, Liu K, Arabzadeh A, et al. SMARCA4 loss increases RNA polymerase II pausing and elevates R-loops to inhibit BRCA1-mediated repair in ovarian cancer. Cancer Res. 2025;85: 2997–3014. doi:10.1158/0008-5472.CAN-24-3990

35. Wagner EJ, Tong L, Adelman K. Integrator is a global promoter-proximal termination complex. Mol Cell. 2023;83: 416–427. doi:10.1016/j.molcel.2022.11.012

36. Battaglia S, Lidschreiber M, Baejen C, Torkler P, Vos SM, Cramer P. RNA-dependent chromatin association of transcription elongation factors and Pol II CTD kinases. Elife. 2017;6. doi:10.7554/eLife.25637

37. Li Z, Li M, Wang D, Hou P, Chen X, Chu S, et al. Post-translational modifications of EZH2 in cancer. Cell Biosci. 2020;10: 143. doi:10.1186/s13578-020-00505-0

38. Yuan H, Han Y, Wang X, Li N, Liu Q, Yin Y, et al. SETD2 Restricts Prostate Cancer Metastasis by Integrating EZH2 and AMPK Signaling Pathways. Cancer Cell. 2020;38: 350–365.e7. doi:10.1016/j.ccell.2020.05.022

39. Zeng Y, Qiu R, Yang Y, Gao T, Zheng Y, Huang W, et al. Regulation of EZH2 by SMYD2-Mediated Lysine Methylation Is Implicated in Tumorigenesis. Cell Rep. 2019;29: 1482–1498.e4. doi:10.1016/j.celrep.2019.10.004

40. Wan J, Zhan J, Li S, Ma J, Xu W, Liu C, et al. PCAF-primed EZH2 acetylation regulates its stability and promotes lung adenocarcinoma progression. Nucleic Acids Res. 2015;43: 3591–3604. doi:10.1093/nar/gkv238

41. Thompson PR, Wang D, Wang L, Fulco M, Pediconi N, Zhang D, et al. Regulation of the p300 HAT domain via a novel activation loop. Nat Struct Mol Biol. 2004;11: 308–315. doi:10.1038/nsmb740

42. Shimoda M, Lyu Y, Wang K-H, Kumar A, Miura H, Meckler JF, et al. KSHV transactivator-derived small peptide traps coactivators to attenuate MYC and inhibits leukemia and lymphoma cell growth. Commun Biol. 2021;4: 1330. doi:10.1038/s42003-021-02853-0

43. García-Nafría J, Watson JF, Greger IH. IVA cloning: A single-tube universal cloning system exploiting bacterial In Vivo Assembly. Sci Rep. 2016;6: 27459. doi:10.1038/srep27459

44. Sanjana NE, Shalem O, Zhang F. Improved vectors and genome-wide libraries for CRISPR screening. Nat Methods. 2014;11: 783–784. doi:10.1038/nmeth.3047

45. Zuris JA, Thompson DB, Shu Y, Guilinger JP, Bessen JL, Hu JH, et al. Cationic lipid-mediated delivery of proteins enables efficient protein-based genome editing in vitro and in vivo. Nat Biotechnol. 2015;33: 73–80. doi:10.1038/nbt.3081

46. Skene PJ, Henikoff JG, Henikoff S. Targeted in situ genome-wide profiling with high efficiency for low cell numbers. Nat Protoc. 2018;13: 1006–1019. doi:10.1038/nprot.2018.015

47. Meers MP, Bryson TD, Henikoff JG, Henikoff S. Improved CUT&RUN chromatin profiling tools. Elife. 2019;8. doi:10.7554/eLife.46314

48. Corces MR, Trevino AE, Hamilton EG, Greenside PG, Sinnott-Armstrong NA, Vesuna S, et al. An improved ATAC-seq protocol reduces background and enables interrogation of frozen tissues. Nat Methods. 2017;14: 959–962. doi:10.1038/nmeth.4396

49. Buenrostro JD, Wu B, Chang HY, Greenleaf WJ. ATAC-seq: A Method for Assaying Chromatin Accessibility Genome-Wide. Curr Protoc Mol Biol. 2015;109: 21.29.1–21.29.9. doi:10.1002/0471142727.mb2129s109

50. Patro R, Duggal G, Love MI, Irizarry RA, Kingsford C. Salmon provides fast and bias-aware quantification of transcript expression. Nat Methods. 2017;14: 417–419. doi:10.1038/nmeth.4197

51. Love MI, Soneson C, Hickey PF, Johnson LK, Pierce NT, Shepherd L, et al. Tximeta: Reference sequence checksums for provenance identification in RNA-seq. PLoS Comput Biol. 2020;16: e1007664. doi:10.1371/journal.pcbi.1007664

52. Love MI, Huber W, Anders S. Moderated estimation of fold change and dispersion for RNA-seq data with DESeq2. Genome Biol. 2014;15: 550. doi:10.1186/s13059-014-0550-8

53. Zhu A, Ibrahim JG, Love MI. Heavy-tailed prior distributions for sequence count data: removing the noise and preserving large differences. Bioinformatics. 2019;35: 2084–2092. doi:10.1093/bioinformatics/bty895

54. Liberzon A, Birger C, Thorvaldsdóttir H, Ghandi M, Mesirov JP, Tamayo P. The Molecular Signatures Database (MSigDB) hallmark gene set collection. Cell Syst. 2015;1: 417–425. doi:10.1016/j.cels.2015.12.004

55. Jalili V, Matteucci M, Masseroli M, Morelli MJ. Using combined evidence from replicates to evaluate ChIP-seq peaks. Bioinformatics. 2018;34: 2338. doi:10.1093/bioinformatics/bty119

56. Yu G, Wang L-G, He Q-Y. ChIPseeker: an R/Bioconductor package for ChIP peak annotation, comparison and visualization. Bioinformatics. 2015;31: 2382–2383. doi:10.1093/bioinformatics/btv145

57. Zhu LJ, Gazin C, Lawson ND, Pagès H, Lin SM, Lapointe DS, et al. ChIPpeakAnno: a Bioconductor package to annotate ChIP-seq and ChIP-chip data. BMC Bioinformatics. 2010;11: 237. doi:10.1186/1471-2105-11-237

58. Lun ATL, Smyth GK. csaw: a Bioconductor package for differential binding analysis of ChIP-seq data using sliding windows. Nucleic Acids Res. 2016;44: e45. doi:10.1093/nar/gkv1191

59. Lun ATL, Smyth GK. De novo detection of differentially bound regions for ChIP-seq data using peaks and windows: controlling error rates correctly. Nucleic Acids Res. 2014;42: e95. doi:10.1093/nar/gku351

60. Heinz S, Benner C, Spann N, Bertolino E, Lin YC, Laslo P, et al. Simple combinations of lineage-determining transcription factors prime cis-regulatory elements required for macrophage and B cell identities. Mol Cell. 2010;38: 576–589. doi:10.1016/j.molcel.2010.05.004

